# SNaQ.jl: Improved Scalability for Phylogenetic Network Inference

**DOI:** 10.1101/2025.11.17.688917

**Authors:** Nathan Kolbow, Sungsik Kong, Tyler Chafin, Joshua Justison, Cécile Ané, Claudia Solís-Lemus

**Affiliations:** Wisconsin Institute for Discovery, University of Wisconsin–Madison, N Orchard St, 53706, WI, USA; Department of Statistics, University of Wisconsin–Madison, University Ave, 53706, WI, USA; Division of Fundamental Mathematical Science, RIKEN Center for Interdisciplinary Theoretical and Mathematical Sciences (iTHEMS), Wako, 351-0198, Saitama, Japan; University of Arkansas, W. Dickson St., 72704, Arkansas, USA; Tree of Life Programme, Wellcome Sanger Institute, Wellcome Trust Genome Campus, CB10 1RQ, Cambridge, United Kingdom; Department of Plant Pathology, University of Wisconsin–Madison, Lincoln Dr, 53706, WI, USA

## Abstract

Phylogenetic networks represent complex biological scenarios that are overlooked in trees, such as hybridization and horizontal gene transfer. Although numerous methods have been developed for phylogenetic network inference, their scalability is severely limited by the computational demands of likelihood optimization and the vastness of network space. Composite (or pseudo-) likelihood approaches like `SNaQ` have improved computational tractability for network inference, but they remain inadequate for datasets of sizes routinely handled by tree inference methods. Here, we introduce `SNaQ.jl`, a new standalone Julia package with the composite likelihood inference originally implemented within `PhyloNetworks.jl` as well as new scalability features that enhance computational efficiency through (1) parallelization of quartet likelihood calculations during composite likelihood computation, (2) weighted random selection of quartets, and (3) probabilistic decision-making during network search. Through a simulation study and empirical data analysis, we show that this new version of `SNaQ.jl` (version 1.1) improves average runtimes by up to 499% on average with no change in function parameters or method accuracy.

## 1 Introduction

Phylogenetic networks depict complex biological scenarios that phylogenetic trees overlook, such as hybrid speciation, introgression, and horizontal gene transfer [Huson and Bryant, 2006, Kong et al., 2025]. Existing network inference methods that utilize full likelihood [Yu et al., 2014, Wen et al., 2016, 2018, Zhang et al., 2018] are not scalable enough for datasets with more than a dozen taxa, due to the high complexity and computational burden of full likelihood equations. To improve scalability, network inference methods that utilize a composite likelihood framework (sometimes referred to as pseudo-likelihood) were introduced. These methods simplify calculations by decomposing the full network into a set of smaller units (typically 4-taxon trees or networks referred to as quartets or quarnets) before computing likelihood calculations on each of these subunits. Then, the overall composite likelihood of the network is obtained as the product of the likelihoods of all quarnets. This approach has been useful in trees (e.g., `MP-EST` [Liu et al., 2010]) and networks (e.g., `SNaQ` [Solís-Lemus and Ané, 2016], `PhyNEST` [Kong et al., 2022b], `PhyloNet` [Yu and Nakhleh, 2015, Zhu and Nakhleh, 2018]) and is much faster than full likelihood methods without compromising method accuracy [Hejase and Liu, 2016]. Despite its computational advantages, composite likelihood estimation remains limited to inference with approximately 30 taxa, which is far below the scale of datasets typically encountered in tree analyses [e.g., One Thousand Plant Transcriptomes Initiative, 2019].

Here, we introduce `SNaQ.jl` v1.1, a Julia package implementing the composite likelihood method for inferring level-1, identifiable, semi-directed phylogenetic networks using gene trees under the multispecies network coalescent model proposed in Solís-Lemus and Ané [2016]. Additionally, we introduce the newest version of this software, version 1.1 (hereafter simply “v1.1”), a new version with improved computational efficiency via (i) parallelization of the quartet composite likelihood calculation, (ii) weighted random selection of quartets, and (iii) probabilistic decision-making during network search. The original implementation of the `SNaQ` method prior to these changes (hereafter simply “v1.0”) is available in `SNaQ.jl` for archival purposes and used here for baseline comparisons. In the following, we briefly introduce the core components of `SNaQ.jl` followed by the key improvements made in v1.1. Then, we present the results of a simulation study comparing the performance of each version before doing the same using an empirical dataset. Our results clearly demonstrate that `SNaQ.jl` significantly improves the computational efficiency of network analyses with no change in method accuracy.

## 2 Methods

### 2.1 A new Julia package: SNaQ.jl

Originally developed as part of the `PhyloNetworks.jl` Julia package [Solís-Lemus et al., 2017], the composite likelihood method `SNaQ` [Species Networks applying Quartets; Solís-Lemus and Ané, 2016] improved scalability of network inference by decomposing the full likelihood into quartet and quarnet likelihoods. In brief, for each set of 4 taxa, (*a, b, c, d*) for instance, there are three possible unrooted quartets that can be observed in a gene tree: 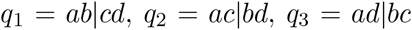. `SnaQ` uses as input the estimated frequency of each of these quartets, denoted as quartet *concordance factors* (CFs) 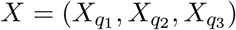 [Baum, 2007] in a set of gene trees across *g* loci.

Next, `SNaQ` searches for the network that best fits the data by traversing the space of topologically identifiable level-1 networks (see Kong et al. [2022a] for a formal definition). Specifically, this means that networks with 2-cycles and most 3-cycles are never proposed during network traversal (for further details see Solís-Lemus and Ané [2016]). In order to quantify how well a given network *N* fits the data, `SNaQ` computes the expected concordance factors for each quarnet *q*_*i*_ according to the multispecies network coalescent model [Yu et al., 2012, Degnan, 2018]. We denote expected concordance factors for a given network *N* as (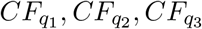). Then, a composite likelihood function is calculated with these data as follows:

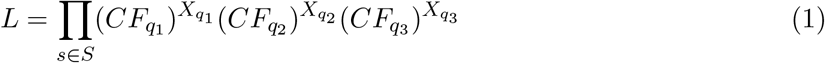

where *S* is the collection of all possible sets of 4 taxa.

Finally, `SNaQ` applies a hill-climbing algorithm to search for the network topology that maximizes the composite likelihood *L*. The algorithm explores the network space using five topological moves: (i) nearest-neighbor interchange (NNI), (ii) addition of a reticulation, (iii) changing the direction of a reticulation edge, and moving either (iv) the target or (v) the origin of an existing reticulation edge. Note that reticulation removal is not considered as a network move because the algorithm uses strict hill-climbing and removal of a reticulation will always result in a lower (or equal) value of *L*.

The network space search begins with an initial network topology *N*_0_ and its corresponding composite likelihood *L*_0_. To explore potential improvements, the algorithm generates a new network topology by randomly applying one of the aforementioned five topological moves. This generates a new network *N*_1_ with composite likelihood *L*_1_. If *L*_1_ *> L*_0_, the new topology is considered an improvement and becomes the focus of subsequent evaluations (i.e., assigning *N*_1_ as *N*_0_). Otherwise, *N*_1_ is discarded and different topological moves are attempted instead. This iterative process continues until no further improvements are found after a predefined number of attempts. Additionally, to mitigate the risk of converging on local optima, `SNaQ` conducts multiple independent runs, by default 10, with varied starting topologies.

### 2.2 Improvements to SNaQ.jl (version 1.1)

#### 2.2.1 Computational parallelizations

`SNaQ.jl` v1.0 parallelizes independent runs across processors, but does not have any parallelization within individual runs. `SNaQ.jl` v1.1 reduces this bottleneck by parallelizing composite likelihood computations and all quartet related operations within individual runs across multiple threads.

#### 2.2.2 Quartet sampling

In `SNaQ.jl` v1.0, every available 4-taxon subset is used when computing the composite likelihood function. This scales quartically with the number of taxa *n* because the total number of 4-taxon sets on *n* taxa is 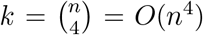. In `SNaQ.jl` v1.1, we introduce the argument `propQuartets`. This argument takes a decimal value in (0, 1] specifying the proportion of 4-taxon subsets that are utilized when computing a network’s composite likelihood. Note that `propQuartets=1` (which remains as the default) corresponds to the approach utilized in v1.0. So, if there are *k* 4-taxon subsets in the CF input table, a subset of ⌈*k* × `propQuartets`⌉ are chosen uniformly at random at the beginning of each run. These subsets are controlled by the initial seed provided to the algorithm, so results are deterministic and reproducible.

This subsampling approach has already been successfully used in phylogenetic inference [Chifman and Kubatko, 2014, 2015]. Once the network topology search is completed, one final round of numerical optimization is performed with all *k* CFs to attain the final network’s full log composite likelihood score.

#### 2.2.3 Topological moves using quartet weighting

In `SNaQ.jl` v1.0, topological moves (iii), (iv), and (v) listed above involve the random selection of a reticulation edge that acts as the focus of the topological modification or “move”. For example, in “move origin”, we select one reticulation node at random and one reticulation edge connected to this reticulation node. Then, the parent or “origin” of this reticulation edge is moved to a randomly chosen neighbor edge. This process is often wasteful as it does not consider whether the current placement of the origin was a good fit for the data.

In `SNaQ.jl` v1.1, we introduce the novel argument `probQR`, which takes a value in [0, 1] and defines the probability of utilizing weighted random selection instead of simple random selection when choosing the reticulation edge at which these topological moves are performed. Note that `probQR=0` (which remains as the default) corresponds to the approach utilized in v1.0. When weighted random sampling is used, each quartet *q*_*i*_ extracted from the current network is given a weight 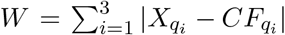 based on the absolute difference between the observed (*X*_*q*_) and the expected (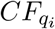) concordance factors. On 4-taxon sets with high weights, the current network CFs fit the data poorly. A 4-taxon set is sampled at random using these weights, and one of the edges spanned by the associated quarnet is sampled uniformly at random to be the focus of the move in order to sample poorly-fitting edges more frequently.

### 2.3 Evaluation using simulated and empirical data

#### 2.3.1 Simulations

To evaluate the performance of `SNaQ.jl` v1.1, we generated a set of species networks with *n* ∈ {10, 20} taxa and *h* ∈ {1, 3} reticulations. To do so, we first generated a species tree under a Yule process using the R package `phytools` [Revell, 2012] for each *n*, then we added *h* reticulations onto the topologies at arbitrary positions. Three of these networks feature primarily shallow reticulations while one features deep reticulations (see Figure S1 for topologies). We ensured that each network was level-1 using the R package `SiPhyNetworks` [Justison et al., 2023]. To simulate low, medium, and high levels of incomplete lineage sorting, we set the average branch length in each network to 2.0, 1.0, and 0.5 respectively, excluding minor hybrid edges which had length 0. Inheritance proportions *γ* were 0.5 in all cases.

Using the Julia package `PhyloCoalSimulations.jl` [Fogg et al., 2023], we generated a set of *g* ∈ {300, 1000, 3000} gene trees for each species network. For each gene tree, we generated a multiple sequence alignment with 1,000 base pairs using `seq-gen` [Rambaut and Grass, 1997], setting the branch length scale parameter to 0.03 and base frequency of nucleotides as *A* = *T* = 0.3 and *C* = *G* = 0.2 under the HKY85 model [Hasegawa et al., 1985]. These sequence alignments were then used to estimate gene trees with `IQ-TREE` (version 1.6.12) [Nguyen et al., 2015] under its automatic model selection option under Bayesian information criterion. Gene tree estimation errors (GTEEs) were computed by dividing the hardwired cluster dissimilarity (HWCD) between each estimated gene tree and its corresponding true gene tree by the maximum possible HWCD value of 2(*n* ™ 3) where *n* is the number of taxa in the tree. Average GTEEs ranged across simulations from 0.21 to 0.32.

Observed CFs were calculated from the estimated gene trees and used as input for `SNaQ.jl` v1.0 and v1.1. Starting topologies were randomly selected from among the estimated gene trees and the true *h* was specified for each simulation. Default optimization parameters were used for both versions of `SNaQ.jl`, but we additionally specified `probQuartets` ∈ {1.0, 0.9, 0.7, 0.5} and `probQR` ∈ {0, 0.5, 1.0} for v1.1. We computed the HWCD between the estimated and true networks using the Julia package `PhyloNetworks.jl` [Solís-Lemus et al., 2017] and recorded the runtime for each network analysis. The analyses were executed with 4, 8, and 16 processors and 2 threads per processor to assess efficiency in different computational settings. This process was repeated 100 times when *n* = 10 and 20 times when *n* = 20 for each set of parameters. For each replicate, new gene trees and multiple sequence alignments were simulated each time. Each replicate analysis was conducted with 10 independent `SNaQ.jl` runs.

Simulations were performed on a computing cluster where hardware is not uniform, but all simulations were limited to a given number of processors and 64-128GB of memory.

#### 2.3.2 Empirical data

We repeated the analysis of 24 swordtails and platyfishes (*Xiphophorus*: Poeciliidae) from Solís-Lemus and Ané [2016] with data consisting of 1,183 gene trees from Cui et al. [2013]. Networks were inferred with *h* = 0 to 5 reticulations where each network with *h >* 0 used the previous best scoring network as its starting point. When *h* = 0, a random estimated gene tree was used as the inference starting point. After obtaining these networks, a single best-fitting network was selected by applying the slope heuristic method implemented in CAPUSHE [Baudry et al., 2012] to the networks’ composite likelihood scores. Inference was performed with 10 processors, 2 threads per processor, and default arguments to illustrate that previous empirical results can be attained much faster as a consequence of the new parallelizations introduced in `SNaQ.jl` v1.1.

## 3 Results and discussion

### 3.1 Simulations

As expected, we observed no noticeable difference in method accuracy between v1.1 and v1.0 when identical parameters were used. Additionally, there was no clear degradation in accuracy as `propQuartets` decreased (see Figures S2-5), even as the number of gene trees and `probQR` were varied.

Next, we highlight improvements in the efficiency of `SNaQ.jl` v1.1 relative to v1.0 (Figure 1 for full runs with 20 taxa and 3 hybridization events, Figures S10-12 for other networks; Figure 2 for individual optimization iteration runtimes). First, `SNaQ.jl` v1.1 utilizes computational resources more efficiently than v1.0. For instance, runtimes for v1.0 changed very little as the number of processors increased, but runtimes for v1.1 fell drastically. This behavior occurred when method parameters were identical between v1.0 and v1.1, so this improvement is the effect of the parallelization across threads.

**Figure 1.**
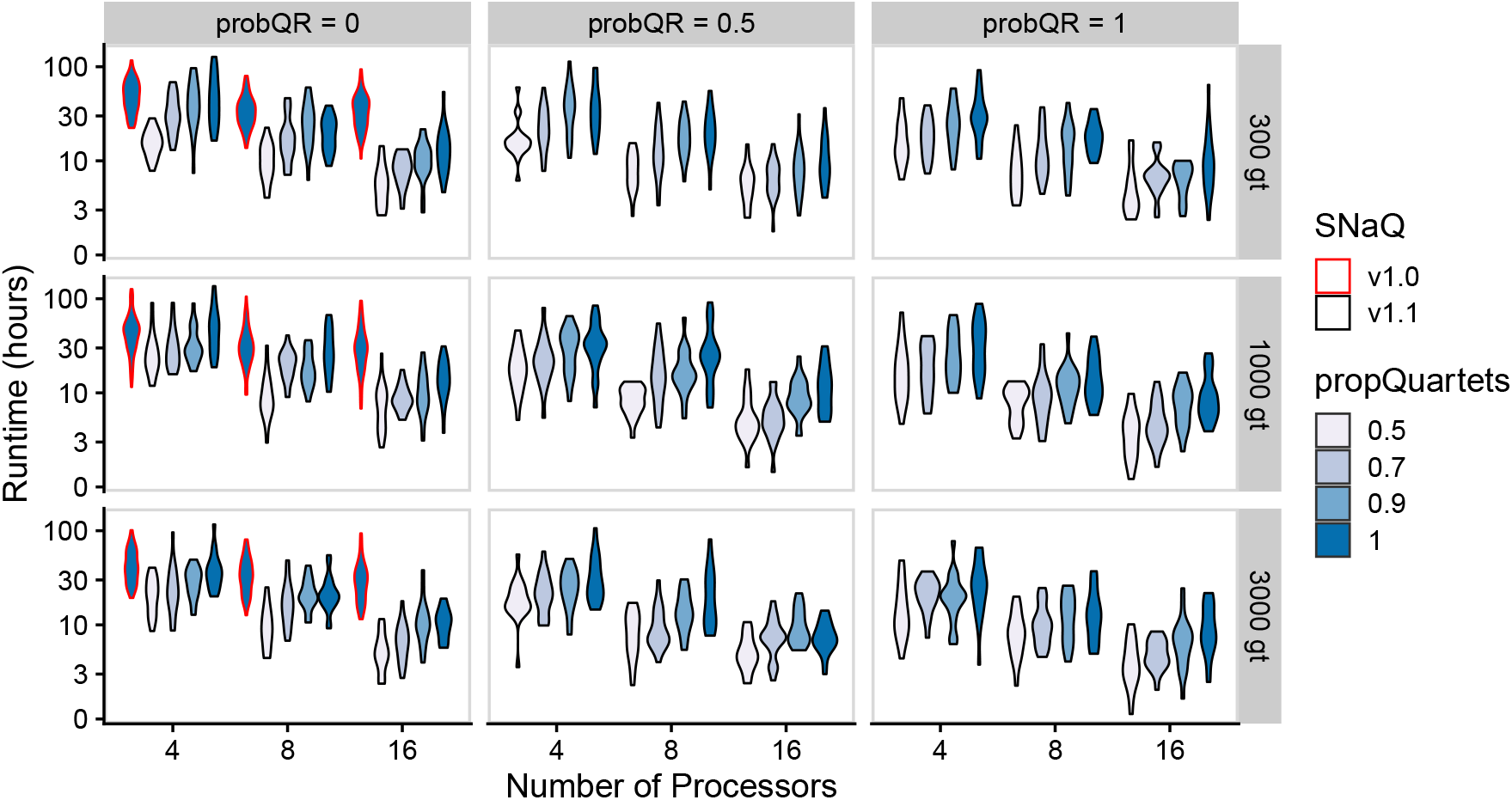
Runtimes (measured in hours and plotted on a log_10_ scale; *y*-axis) for the network with 20 taxa and 3 reticulations. `probQR`, the number of input gene trees (gt), and the number of processors used was varied across simulations. `SNaQ.jl` v1.0 results are outlined in red while v1.1 results are outlined in black. Each violin plot covers 20 replicates.

**Figure 2.**
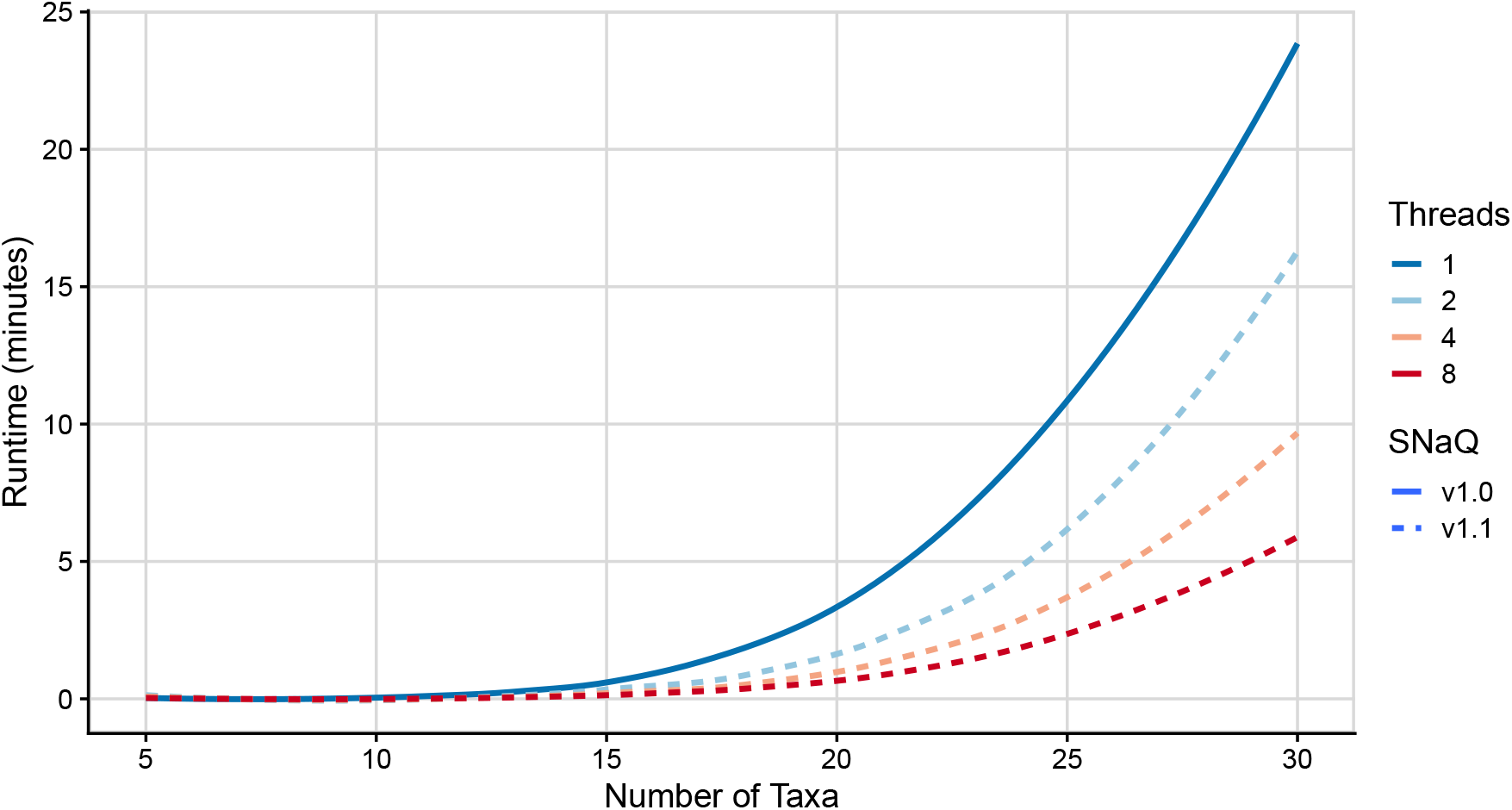
Runtimes (measured in minutes; *y*-axis) taken to optimize the parameters of individual networks using `SNaQ.jl` v1.0 and v1.1. v1.1 was run with 2, 4, and 8 threads, while v1.0 was only run with 1 thread. Results are averaged across replicates with 0-3 hybridizations.

While we expected higher `probQR` values to reduce the time spent searching for networks or to improve inference accuracy, there was no noticeable improvement in either metric when `probQR` was changed. We suspect that higher values of `probQR` could be reducing the overall stochasticity of the search algorithm resulting in higher chances of being stuck in local optima. On the other hand, alternative weighting methods or different network topologies could yield different results but are outside the scope of this study. Such adaptations would be interesting to examine in future work. The effect of `propQuartets` on the performance of `SNaQ.jl` v1.1 was clear in our simulations. Decreasing `propQuartets` led to significant improvements in runtime performance without no noticeable drop in accuracy. The most significant average speed improvement observed in these simulations was by a factor of 811% on the network with *n* = 20, *h* = 1 with 16 processors, *g* = 1000, `probQR`= 1, and `propQuartets`= 0.5. Additionally, across all simulations with 16 processors, v1.1 cuts runtimes compared to v1.0 by factors of 499%, 576%, and 757% with `propQuartets` = 1.0, 0.9, and 0.7 respectively.

When `propQuartets` *<* 1, a simple random sampling approach is taken, but a more informative sampling approach, such as the weighted sampling approach of [Zhang and Mirarab, 2022], may lead to more robust results and enable even smaller values of `propQuartets` to be utilized while retaining overall method accuracy.

Lastly, the runtime improvements seen in parameter optimizations on individual networks (Figure 2) are roughly proportional with respect to the number of threads used, indicating an efficient multi-threading implementation. It is clear, though, that while these improvements drastically improve the time taken to optimize a given network’s parameters, they do not significantly extend the size of network that can be efficiently inferred. This is especially evident when considering that larger networks naturally have larger spaces to traverse, warranting the use of more time consuming optimization routines. Thus, researchers who wish to infer drastically larger networks should continue to use more appropriate methods such as `InPhyNet` [Kolbow et al., 2025] when the primary concern is the biological accuracy of the inferred network, or `Neighbor-Net` [Bryant and Moulton, 2004] when the primary concern is in understanding the discordance present in the set of inferred gene trees.

### 3.2 Empirical data

`SNaQ.jl` requires the number of hybridizations in the inferred network to be specified prior to inference. To pick the best-fitting network to the observed data, networks were inferred with *h* = 0 to 5 reticulations and default function arguments, after which a single best-fitting network was selected by applying the model selection method implemented in the CAPUSHE R package [Baudry et al., 2012] to the networks’ composite likelihood scores. Then, we inferred an additional network with this selected value (*h* = 2) and `propQuartets` = 0.1, 0.3, 0.5, 0.7 to assess how empirical conclusions may differ when utilizing less quartet data.

`SNaQ.jl` v1.1 with default function arguments took 54 hours in aggregate to compute all 6 network topologies (each of which has different *h*), which is 346.2 fewer hours (14.4 fewer days) than v1.0 took originally, equating to a runtime improvement by a factor of 641%. The longest runs for v1.0 and v1.1 were 208.8 hours and 16.5 hours, respectively, equating to an improvement in worst-case runtimes by a factor of 1165%. `SNaQ.jl` v1.1 with `propQuartets` = 0.1, 0.3, 0.5, 0.7 and *h* = 2 took 3.6, 8.2, 6.6, and 17.35 hours to infer, respectively, equating to runtime improvements by factors of 339%, 93%, 139%, and -9% compared to v1.1 with `propQuartets` = 1.0, respectively. Next, we examine topological differences in the networks inferred by v1.0 and v1.1 (see Figures S13-S18 in the Supplementary Material). Networks with *h* = 0 were identical when inferred with both versions and were also identical to the total evidence tree in Cui et al. [2013]. Networks with *h* = 1 contained the same hybridization event and differed only in a small change in the placement of *X. milleri*. Both networks with *h* = 2 identified this hybridization as well. Additionally, they identified the same target for their second hybridization (*X. helleri*) but slightly different origins (*X. clemenciae* versus its ancestor). Similarly, networks with *h* = 3 and *h* = 4 contained these two identical hybridizations but differed in the placement of their remaining hybridizations. The difference in composite log likelihood scores between v1.0 and v1.1 for *h* = 2, 3, 4 were each small (≤ 3%). Finally, networks with *h* = 5 had the same inferred topologies. Inferred inheritance probabilities (*γ*) were roughly similar between the two versions, regardless of differences in the placements of hybridizations.

Despite the similarity of these networks to those in Solís-Lemus and Ané [2016], the network inferred by `SNaQ.jl` v1.1 with *h* = 1 had a drastically better composite log likelihood score than its counterpart inferred by `SNaQ.jl` v1.0. As a result, the best-fitting network determined by slope heuristics would be the network inferred with *h* = 1, in contrast to Solís-Lemus and Ané [2016], where the network with *h* = 2 was selected.

Finally, we denote the best-fitting network inferred with *h* = 2 and `propQuartets` = *p* as 𝒩_*p*_. Networks 𝒩_0.1_ and 𝒩_0.7_ had similar composite log likelihood scores to 𝒩_1.0_ while 𝒩_0.3_ and 𝒩_0.5_ had *better* scores than 𝒩_1.0_ (see Figure S20). Additionally, the topologies of these networks were all remarkably similar. Every network had the same major tree and the same reticulation from *X. xiphidium* to the northern platyfish clade. Further, the second reticulation in each network was always near the siblings *X. monticolus* and *X. clemenciae*, as well as the species *X. helleri*, but the exact location and direction of the reticulation varied.

The networks with 𝒩_0.3_ and 𝒩_0.5_, which had the best composite log likelihood score, estimated inheritance parameters of *γ* = 0.43 and *γ* = 0.32, whereas the other networks estimated *γ* = 0.028 and *γ* = 0.029. These differences stand out because they lead to drastically different interpretations. Most strikingly was that 𝒩_0.1_ was *identical* to 𝒩_1.0_. We assumed that very low values of `propQuartets` would naturally lead to poorly fitting results, which is why our simulation study only went down to `propQuartets` = 0.5, but this result clearly disproves that notion. This result is surprising, but supported by Rosas-Puchuri et al. [2025], who use even fewer quartets and still achieve high levels of inference accuracy (see Figure 5 in [Rosas-Puchuri et al., 2025]).

## 4 Conclusion

`SNaQ.jl` v1.1 implements parallel computing techniques to significantly improve its computational efficiency when run on a machine with multiple processing cores and threads. In our simulation study, we observed runtime improvements by 499% when comparing v1.1 to v1.0 when identical arguments were utilized with 16 processors and 2 threads per processor. Further, we observed average runtime improvements by 757% when the new optimization parameter `propQuartets` was reduced to 0.5. These runtime improvements come without any significant change in accuracy, indicating the v1.1 should be preferred over v1.0. This efficiency was further substantiated by re-analyzing an empirical *Xiphophorus* fish dataset with 24 taxa where runtimes improved by a similar magnitude and major findings in the original analysis were confirmed.

In addition, `SNaQ.jl` v1.1 implements two new parameters to the `snaq!` function: `propQuartets` and `probQR`. Our simulation study shows that reducing `propQuartets` as low as 0.5 significantly improves runtime with no significant difference in accuracy. Further, our empirical results recover identical networks when using `propQuartets` = 1.0 and `propQuartets` = 0.1, suggesting that even smaller subsets of quartets can lead to useful results. These results may be partially dependent on the specific network topologies analysed here, though, so further investigation is warranted to verify the robustness of this parameter to various topological features. Additionally, future work could build on these results by identifying values of `propQuartets` for which `SNaQ.jl`’s accuracy begins to deteriorate, or by improving upon the current implementation by utilizing more sophisticated sampling strategies. On the other hand, `probQR` appeared to have no effect on method accuracy or runtime, so further investigation is warranted to assess why this is the case, and whether different weighted sampling approaches could yield better results.

## 5 Code and data availability

`SNaQ.jl` is a new open source Julia package available at https://github.com/JuliaPhylo/SNaQ. jl. Scripts for simulation and empirical data analyses can be found on Zenodo (https://doi.org/10.5281/zenodo.18664095). Original data used and generated in these simulations can be found on Dryad (https://doi.org/10.5061/dryad.pnvx0k721).

## 6 Competing interests

No competing interest is declared.

## 7 Author contributions statement

CSL and TC conceived the idea for the scalability implementations. TC implemented the improvements into `SNaQ.jl` version 1.1. NK and SK designed the simulations and prepared the manuscript. NK conducted the empirical and simulated analyses. JJ, NK, CA, and CSL refactored the original codebase, split it into two modular packages (`PhyloNetworks.jl` and `SNaQ.jl`), implemented unit tests, and wrote documentation.

## 8 Acknowledgments

This work was supported by the National Science Foundation (DEB-2144367 to CSL and DBI-2010774 to TC).

## Appendix

### A1 Simulation Study

#### A1.1 Simulated Network Topologies

Below are the network topologies used in the simulation study. Note that in each simulation, branch lengths were adjusted as described in the main text to simulate varying levels of incomplete lineage sorting. Specifically, to simulate low, medium, and high levels of ILS, we multiply every edge in the network by a constant such that the average branch length in the network was 2.0, 1.0, or 0.5 respectively.

**Figure 3.**
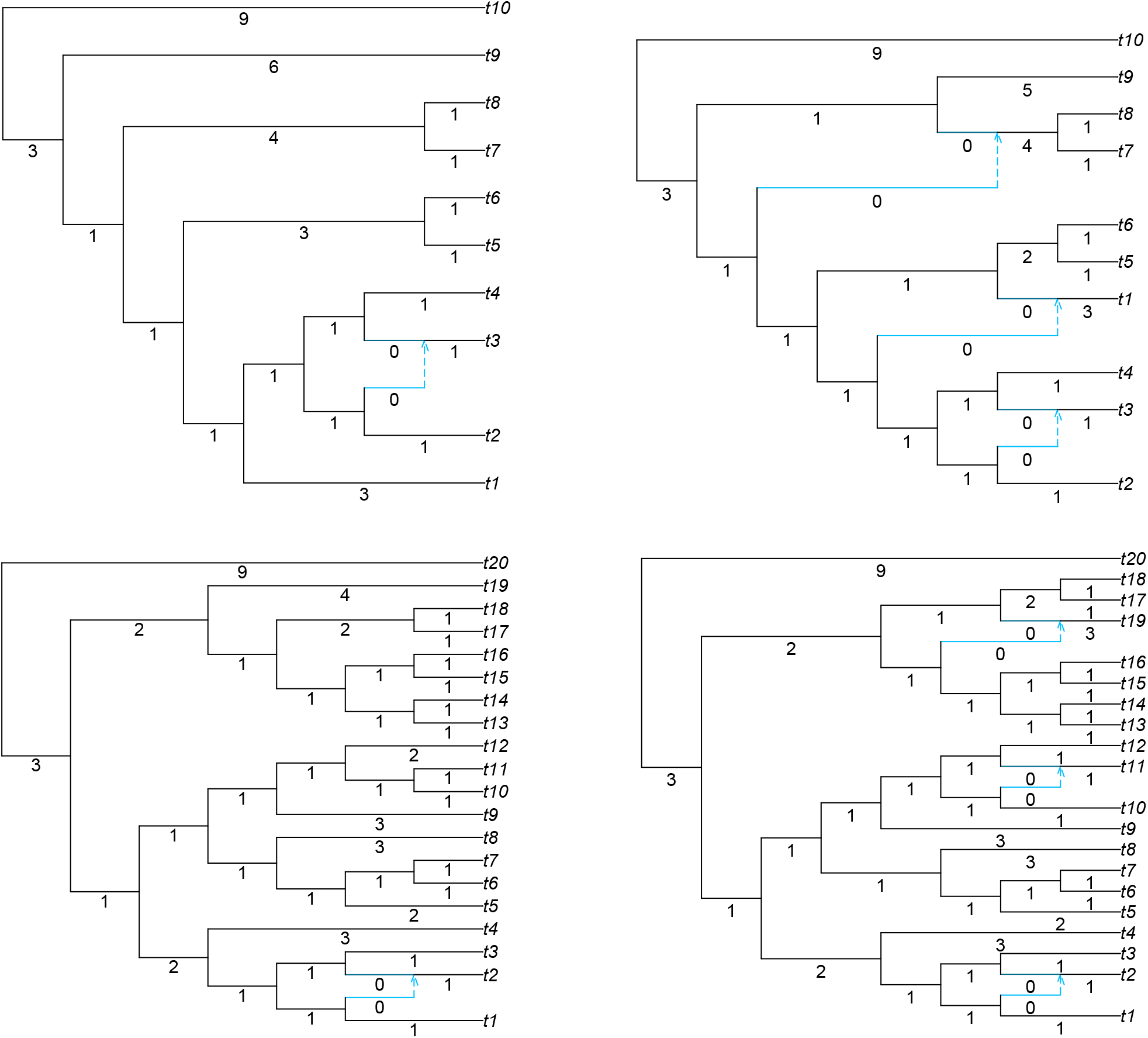
Topologies and edge lengths of all networks used in the simulation study. All inheritance proportions (*γ*, not shown) are 0.5 in each network.

#### A1.2 Accuracy results

**Figure 4.**
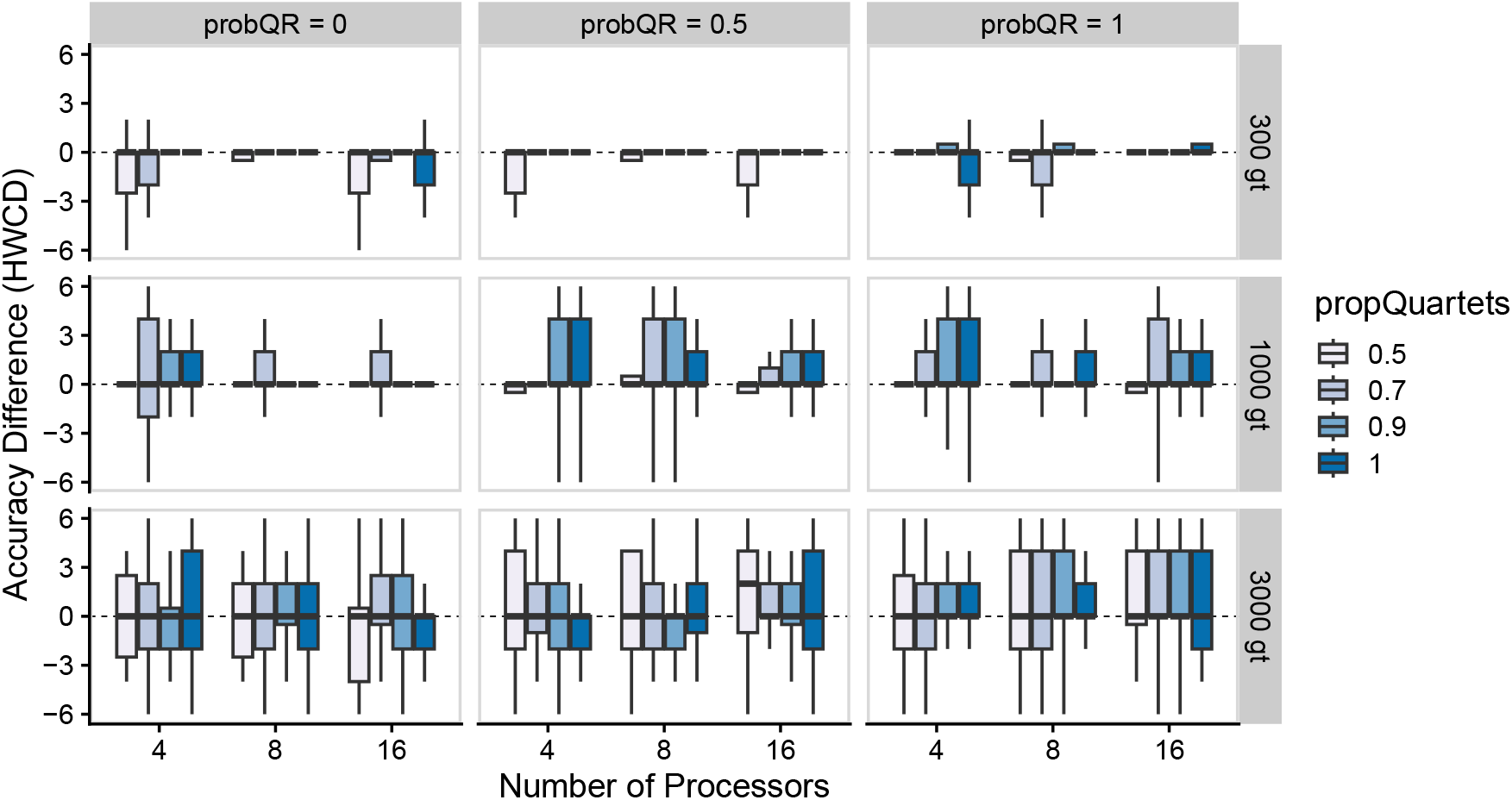
Differences in the accuracies of networks inferred with 10 taxa and 1 reticulation between `SNaQ.jl` v1.0 and v1.1 measured in hardwired cluster distance (HWCD) across an array of simulation parameters. Positive numbers represent v1.1 inferring a more accurate network than v1.0, while negative numbers represent the inverse, and a value of 0 represents networks with equal accuracy relative to the ground truth.

**Figure 5.**
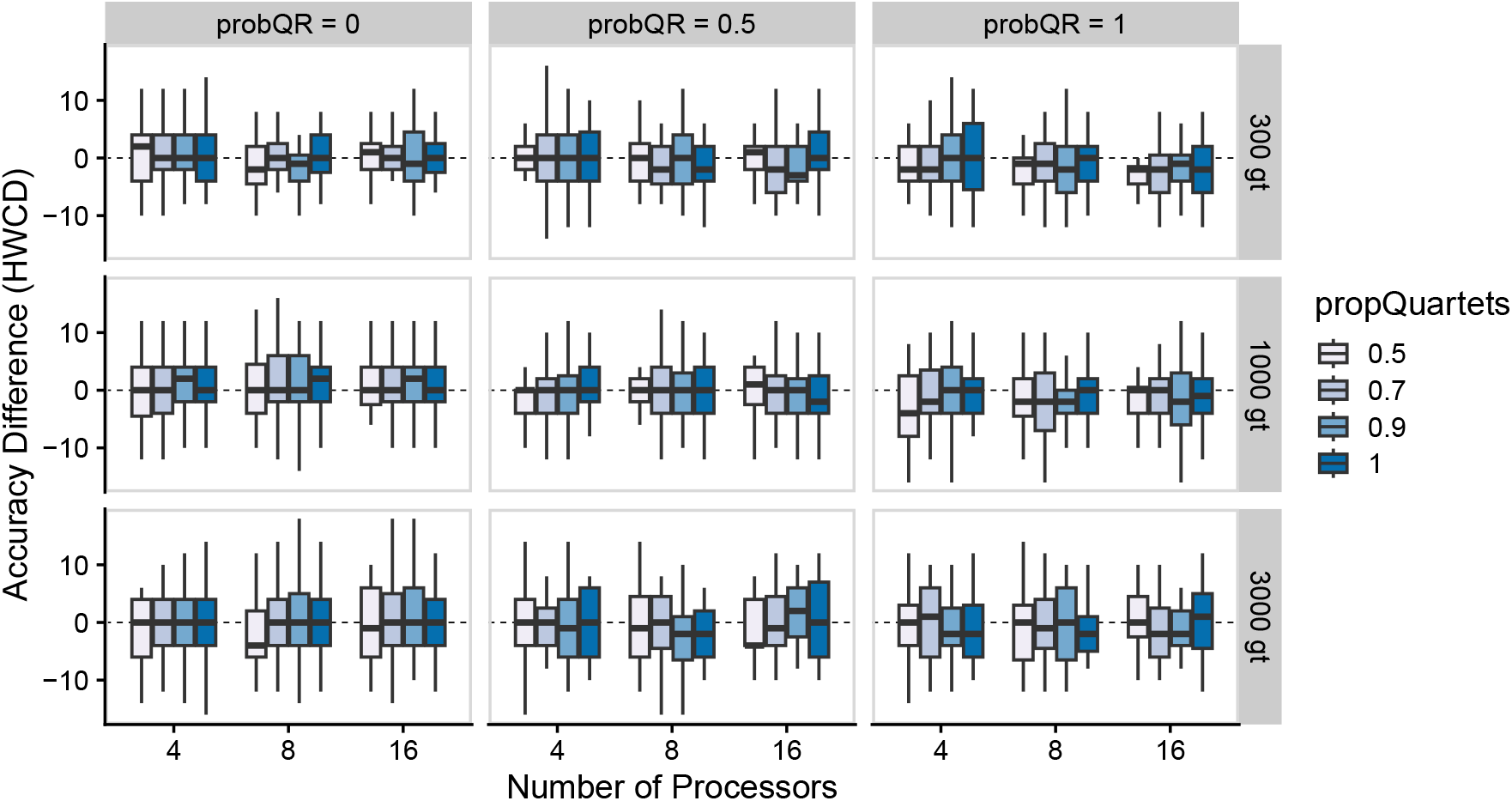
Accuracy results in hardwired cluster distance (HWCD) for the topology with 10 taxa and 3 reticulations across a wide array of simulation parameters. Violin plots outlined in red represent results for `SNaQ.jl` version 1.0, whereas all others represent results for version 1.1.

**Figure 6.**
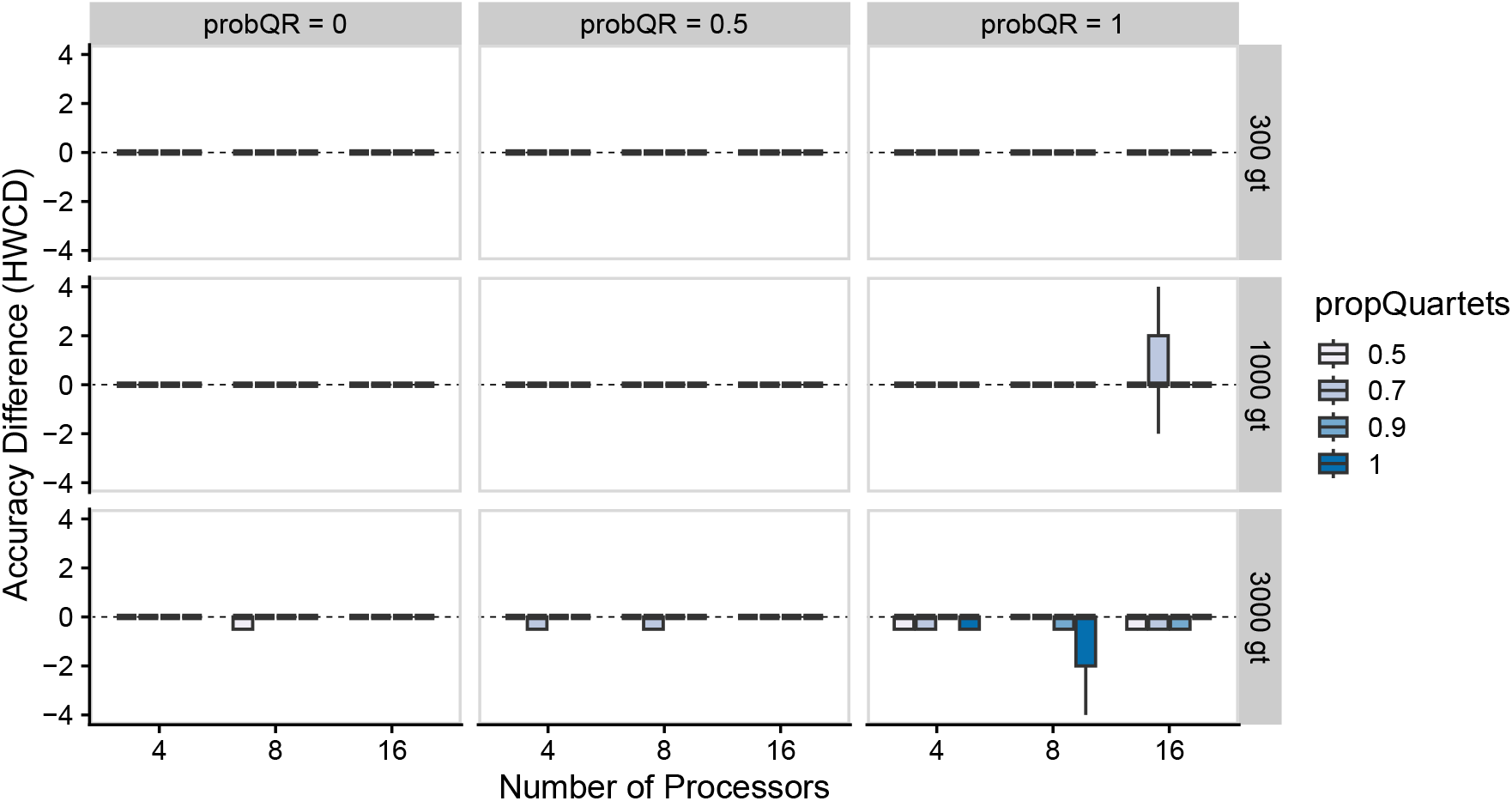
Differences in the accuracies of networks inferred with 20 taxa and 1 reticulation between `SNaQ.jl` v1.0 and v1.1 measured in hardwired cluster distance (HWCD) across an array of simulation parameters. Positive numbers represent v1.1 inferring a more accurate network than v1.0, while negative numbers represent the inverse, and a value of 0 represents networks with equal accuracy relative to the ground truth. This network was particularly easy for `SNaQ.jl` v1.0 and v1.1 to achieve a HWCD value of exactly 4, but very difficult to improve past that point, which is why most values here are exactly 0 (see also Fig 10).

**Figure 7.**
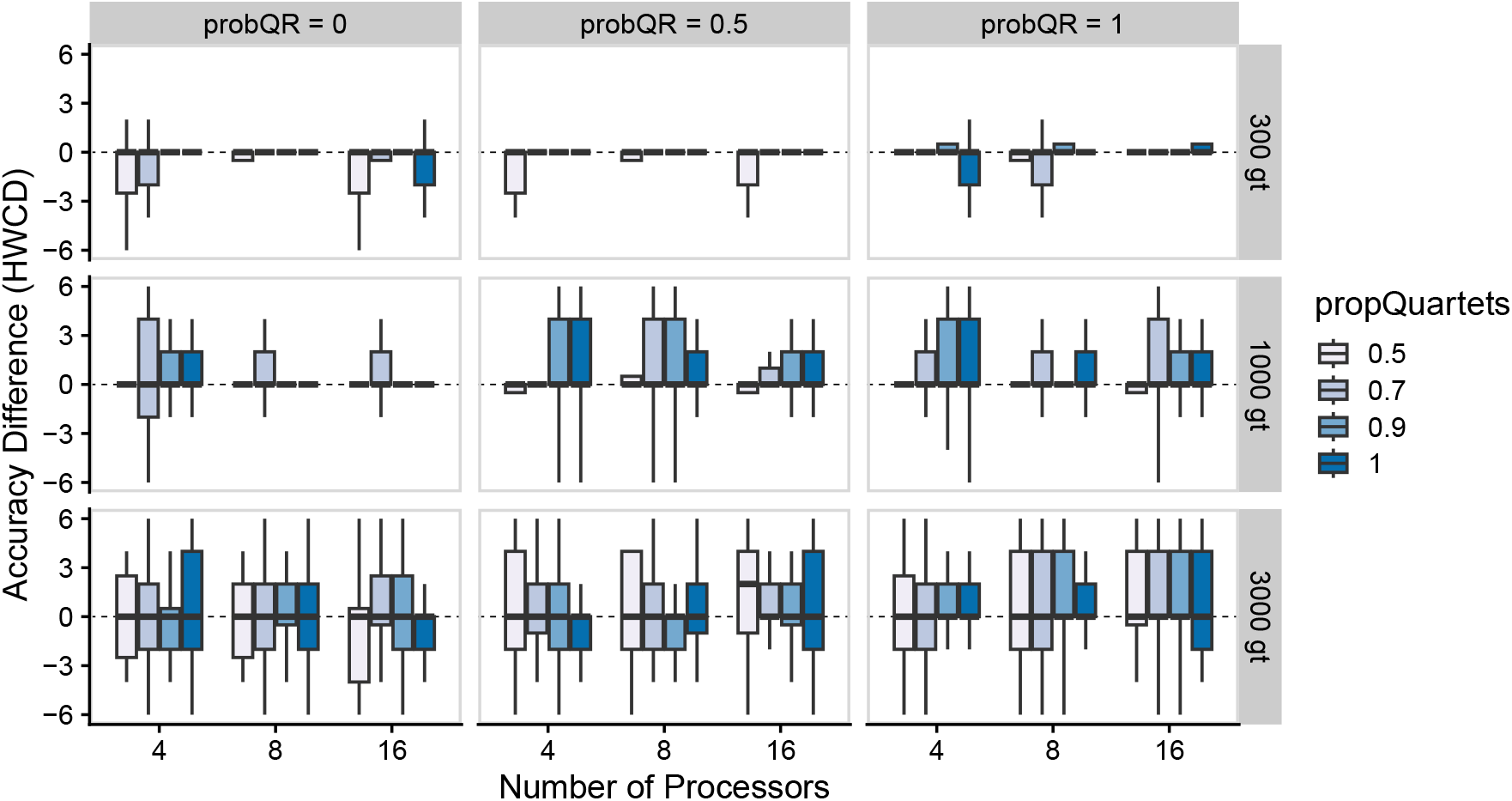
Differences in the accuracies of networks inferred with 20 taxa and 3 reticulation between `SNaQ.jl` v1.0 and v1.1 measured in hardwired cluster distance (HWCD) across an array of simulation parameters. Positive numbers represent v1.1 inferring a more accurate network than v1.0, while negative numbers represent the inverse, and a value of 0 represents networks with equal accuracy relative to the ground truth.

**Figure 8.**
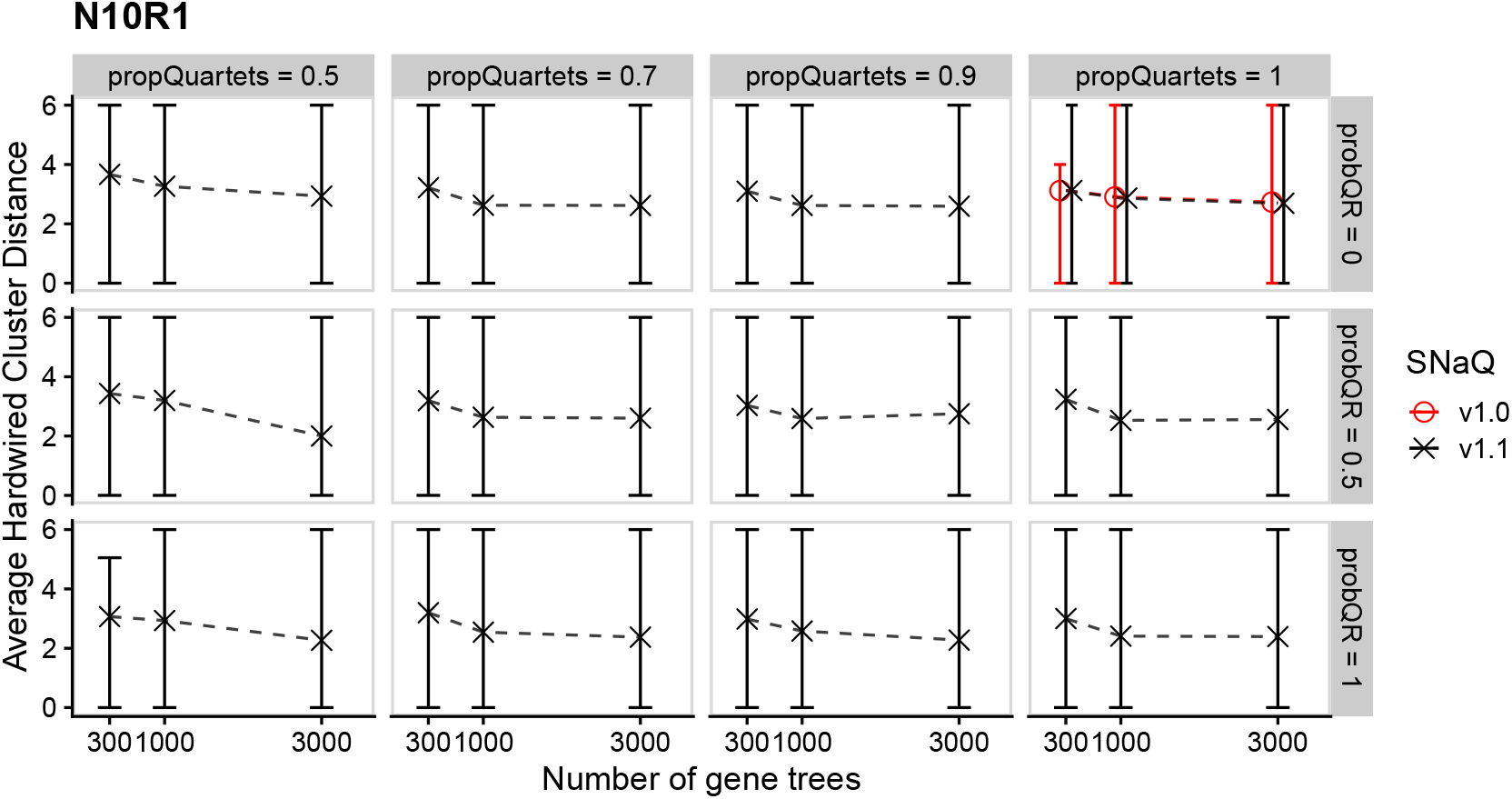
Accuracy results in hardwired cluster distance (HWCD) for the topology with 10 taxa and 1 reticulation relative to the number of input gene trees across a wide array of simulation parameters. Margins represent 95% empirical confidence intervals for observed accuracies, while point estimates are the mean HWCDs for each parameter combination. Simulations utilizing `SNaQ.jl` version 1.0 are shown in red, whereas those utilizing `SNaQ.jl` version 1.1 are shown in blue.

**Figure 9.**
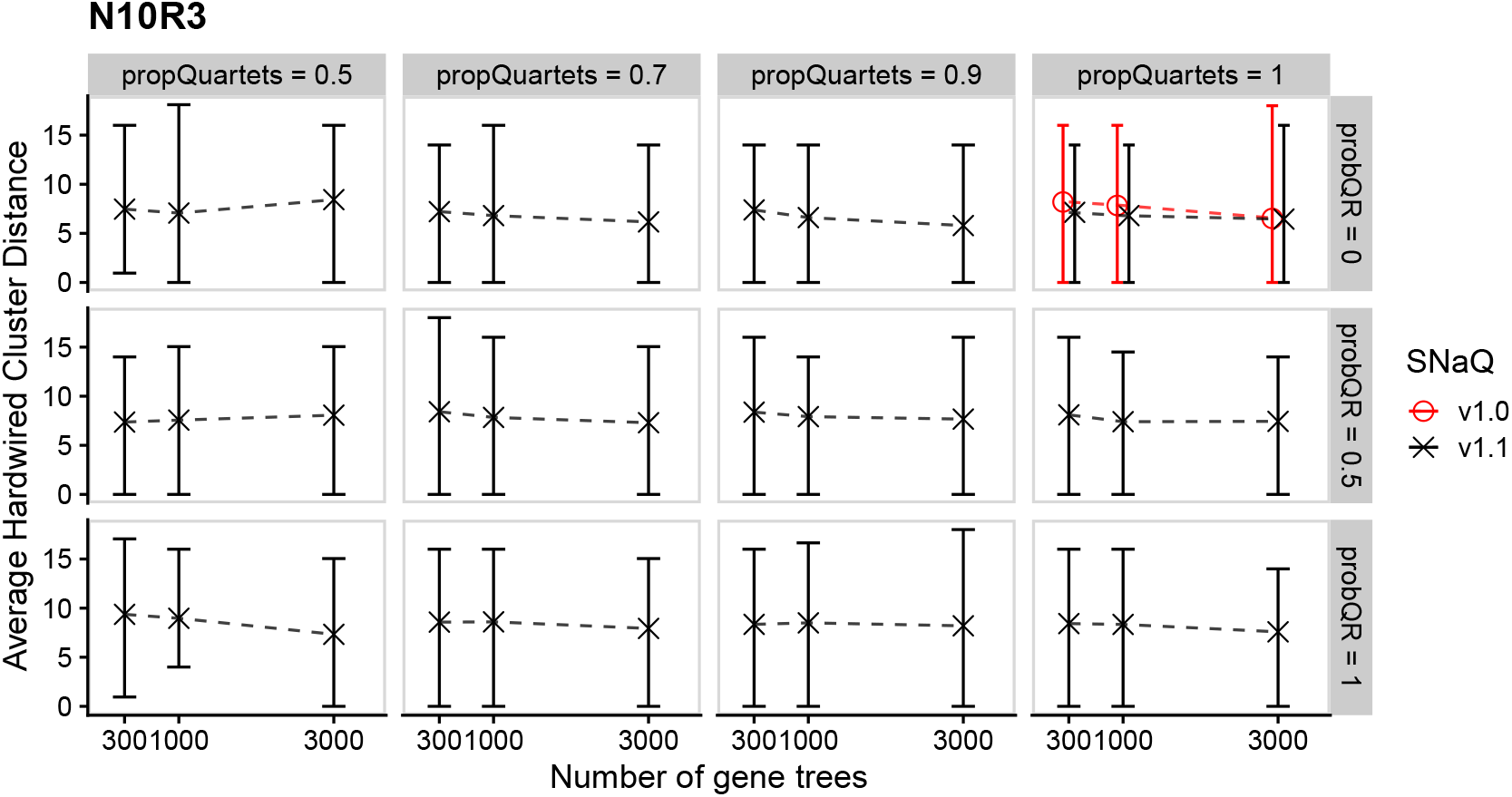
Accuracy results in hardwired cluster distance (HWCD) for the topology with 10 taxa and 3 reticulations relative to the number of input gene trees across a wide array of simulation parameters. Margins represent 95% empirical confidence intervals for observed accuracies, while point estimates are the mean HWCDs for each parameter combination. Simulations utilizing `SNaQ.jl` version 1.0 are shown in red, whereas those utilizing `SNaQ.jl` version 1.1 are shown in blue.

**Figure 10.**
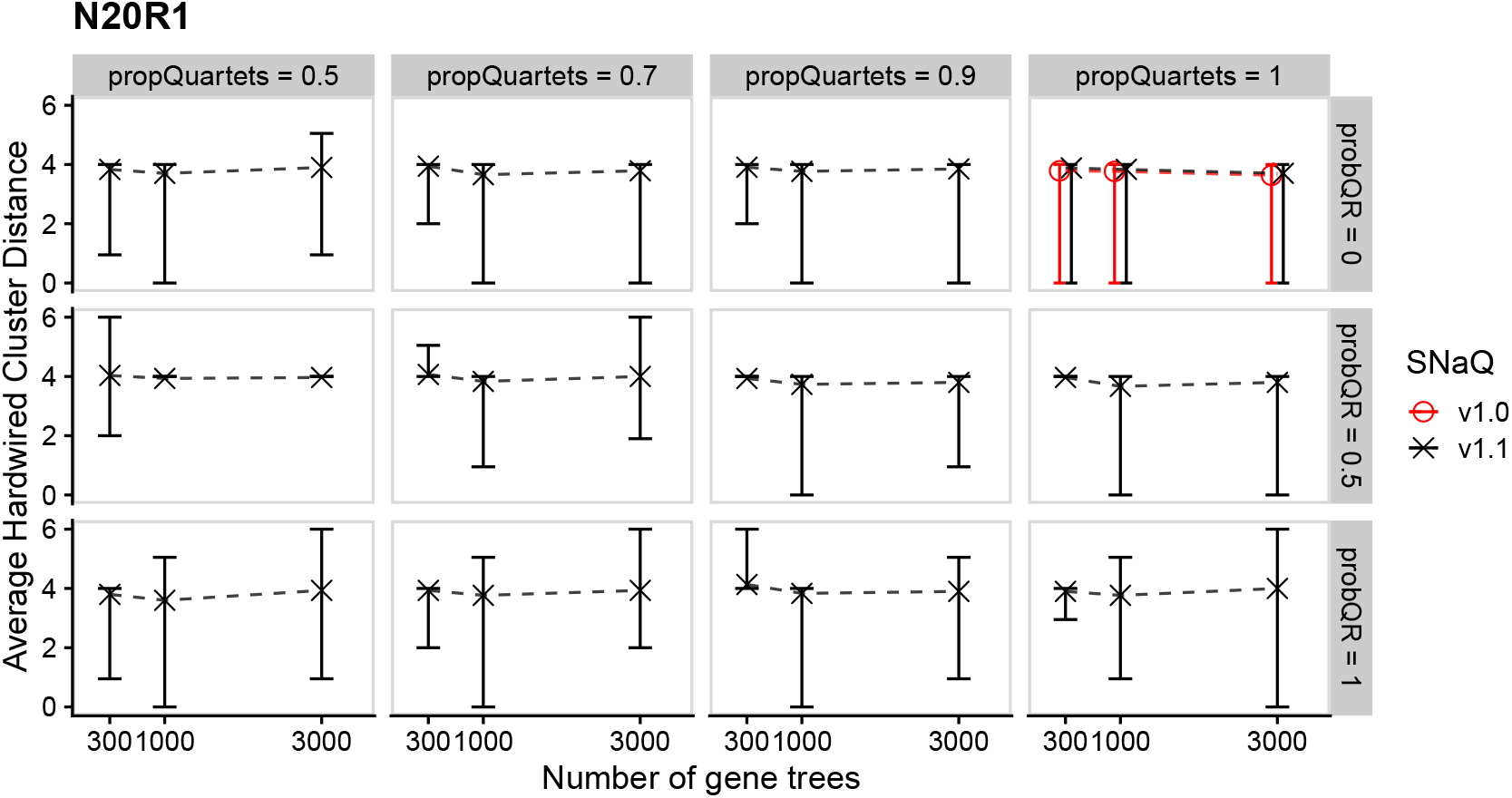
Accuracy results in hardwired cluster distance (HWCD) for the topology with 20 taxa and 1 reticulation relative to the number of input gene trees across a wide array of simulation parameters. Margins represent 95% empirical confidence intervals for observed accuracies, while point estimates are the mean HWCDs for each parameter combination. Simulations utilizing `SNaQ.jl` version 1.0 are shown in red, whereas those utilizing `SNaQ.jl` version 1.1 are shown in blue.

**Figure 11.**
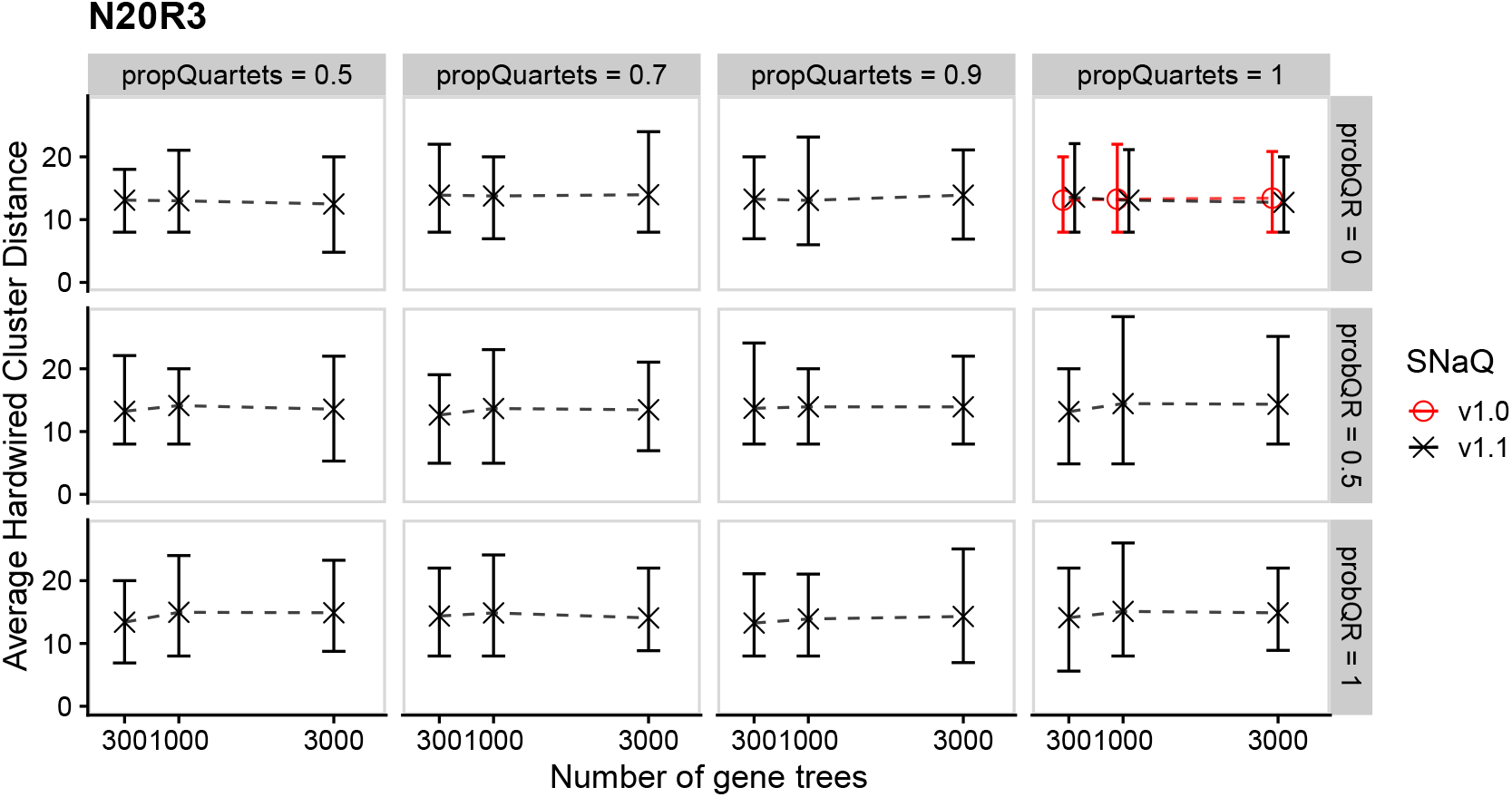
Accuracy results in hardwired cluster distance (HWCD) for the topology with 20 taxa and 3 reticulations relative to the number of input gene trees across a wide array of simulation parameters. Margins represent 95% empirical confidence intervals for observed accuracies, while point estimates are the mean HWCDs for each parameter combination. Simulations utilizing `SNaQ.jl` version 1.0 are shown in red, whereas those utilizing `SNaQ.jl` version 1.1 are shown in blue.

#### A1.3 Runtime results

**Figure 12.**
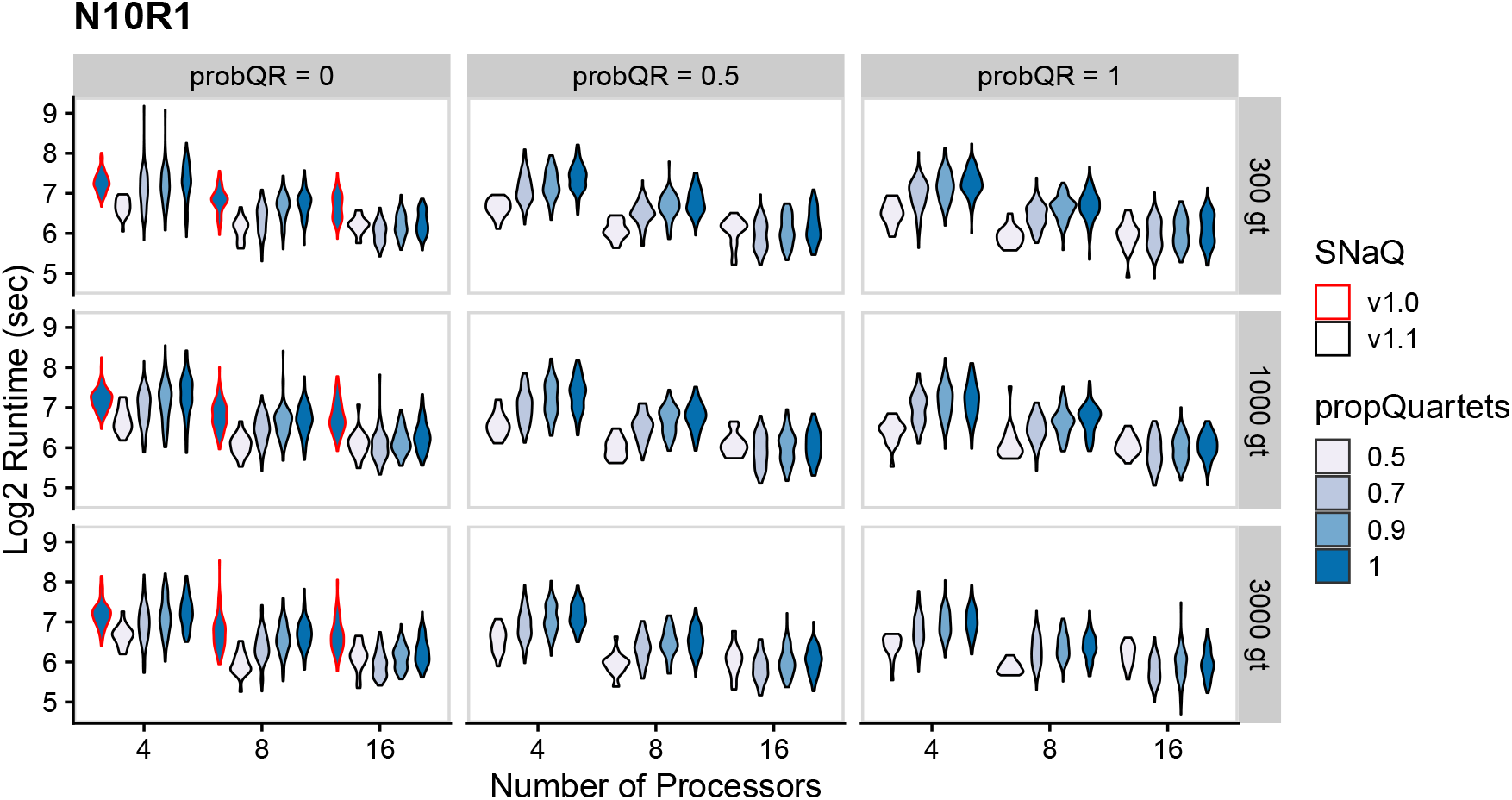
Log2 runtime results in seconds for the topology with 10 taxa and 1 reticulation across a wide array of simulation parameters. Violin plots outlined in red represent results for `SNaQ.jl` version 1.0, whereas all others represent results for version 1.1.

**Figure 13.**
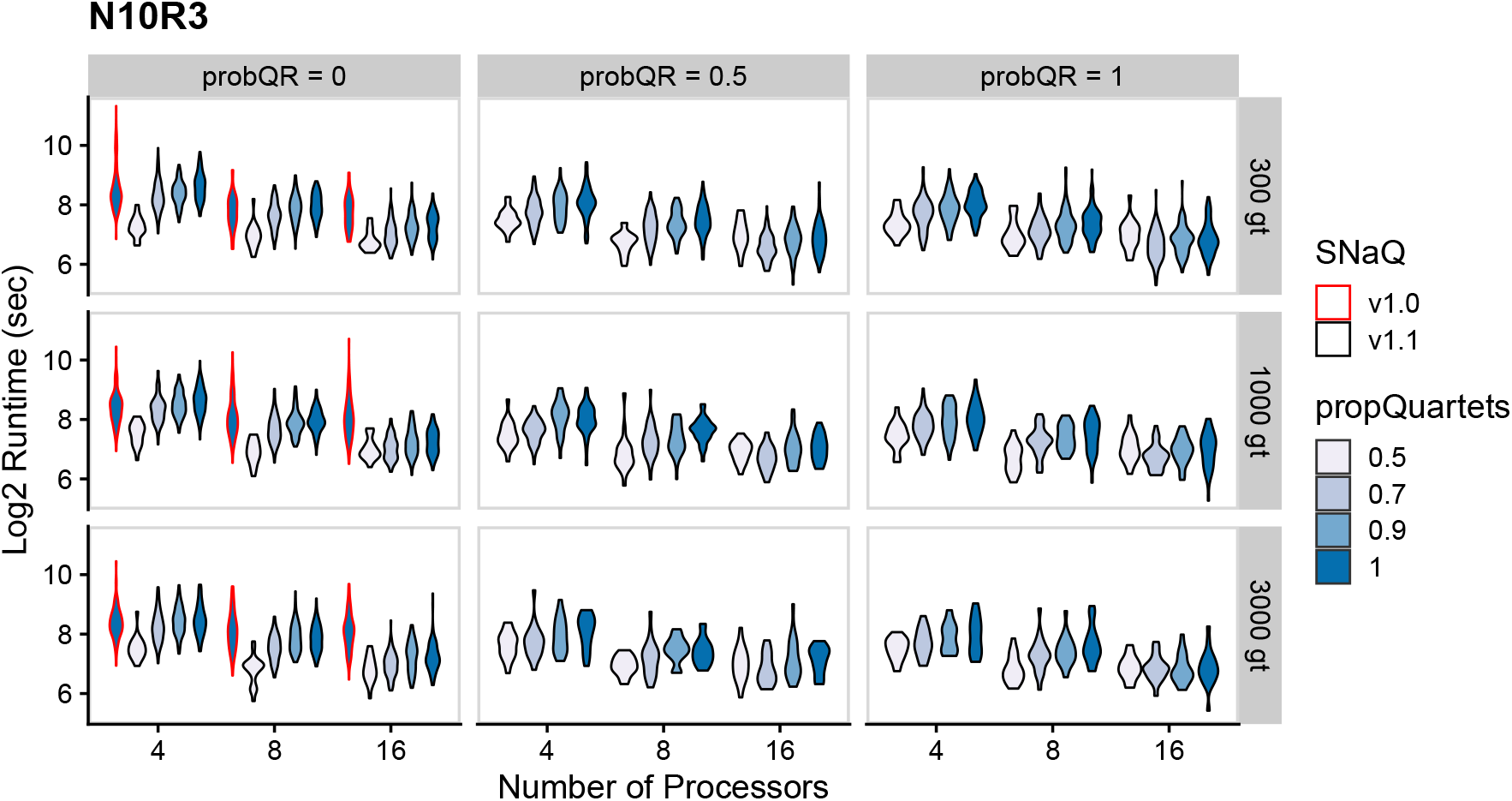
Log2 runtime results in seconds for the topology with 10 taxa and 3 reticulations across a wide array of simulation parameters. Violin plots outlined in red represent results for `SNaQ.jl` version 1.0, whereas all others represent results for version 1.1.

**Figure 14.**
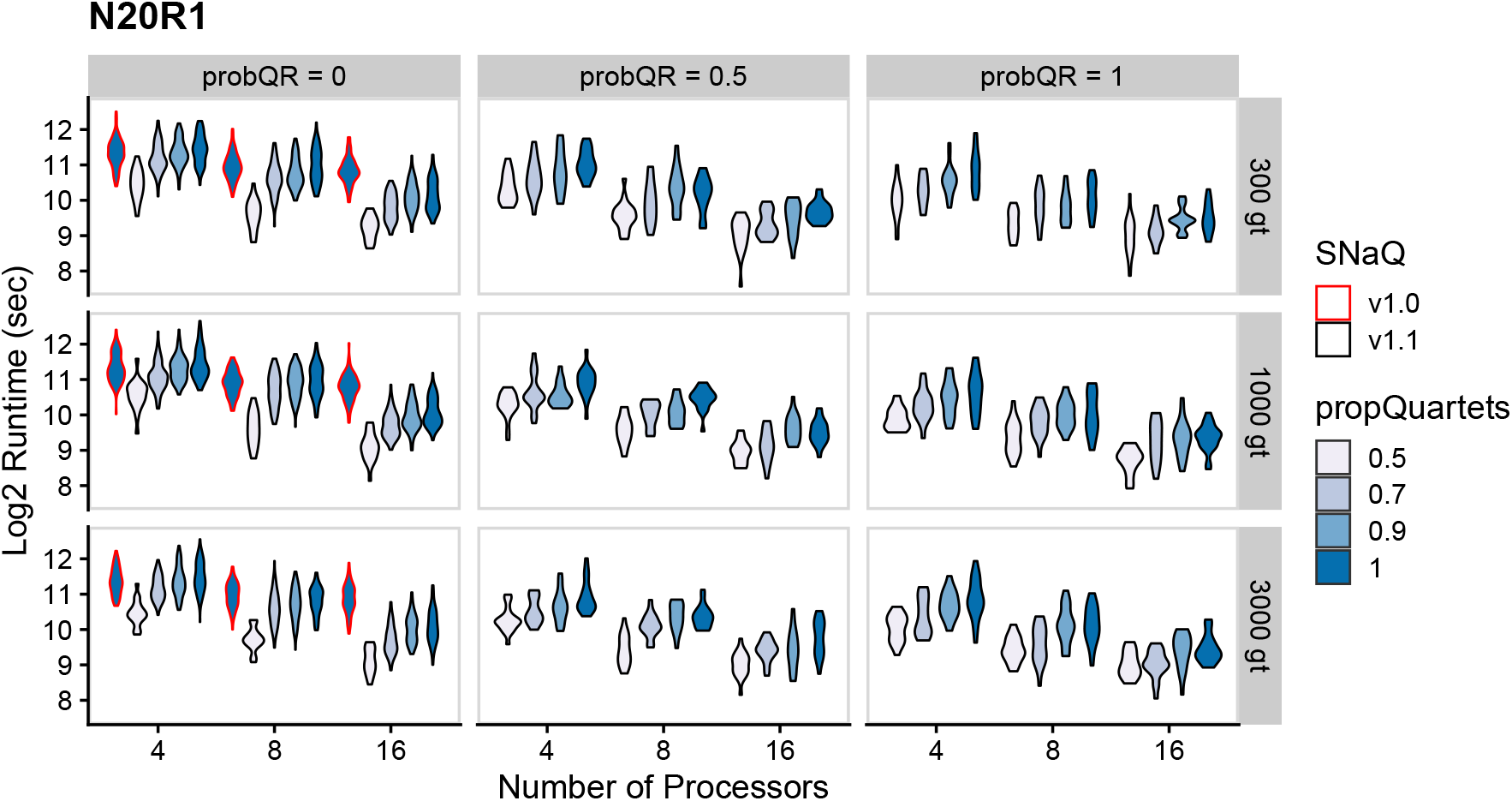
Log2 runtime results in seconds for the topology with 20 taxa and 1 reticulation across a wide array of simulation parameters. Violin plots outlined in red represent results for `SNaQ.jl` version 1.0, whereas all others represent results for version 1.1.

### A2 Empirical Results

**Table 1:** Runtimes (measured in hours) and improvements from SNaQ.jl v1.0 to v1.1 on empirical data with 24 taxa and networks inferred with *h* reticulations using 10 CPU cores and 2 threads per core. The difference in efficiency measured in hours is also provided.

| Reticulation ($h$) | Runtimes (hours) | | | | | |
| --- | --- | --- | --- | --- | --- | --- |
|  | 0 | 1 | 2 | 3 | 4 | 5 |
| **Version 1.0** | 17.3 | 29.3 | 37.8 | 58.2 | 208.8 | 48.8 |
| **Version 1.1** | 8.0 | 16.5 | 15.8 | 3.2 | 4.4 | 6.1 |
| **Difference** | 9.3 | 12.9 | 22.0 | 55.1 | 204.3 | 42.7 |

#### A2.1 Inferred networks

Below are all networks topologies inferred for species *Xiphophorus*: Poeciliidae with *h* ∈ {0, 1, 2, 3, 4, 5} inferred by `SNaQ.jl` versions 1.0 and 1.1 with *X. mayae* arbitrarily chosen as outgroup in all figures.

**Figure 15.**
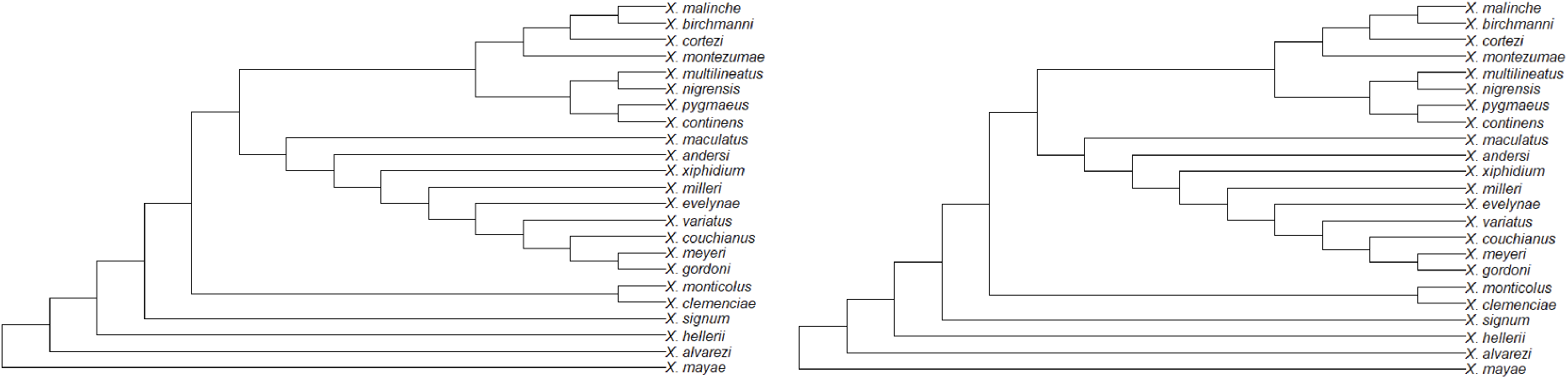
Highest negative log likelihood empirical network inferred by `SNaQ.jl` version 1.1 (left; negative log composite likelihood ≈ 12306) and version 1.0 (right; negative log composite likelihood ≈ 12306) with 0 hybridizations.

**Figure 16.**
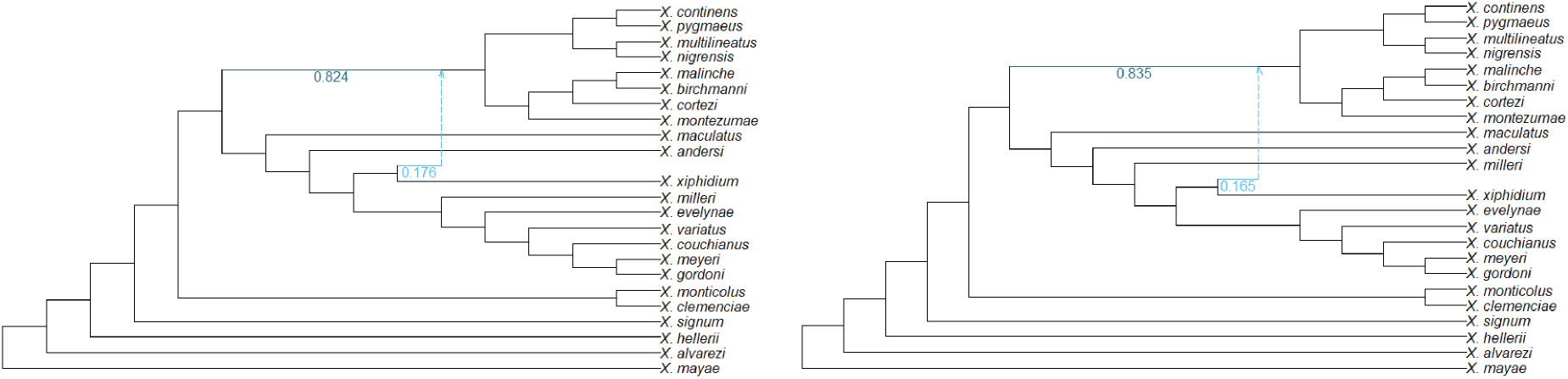
Highest negative log likelihood empirical network inferred by `SNaQ.jl` version 1.1 (left; negative log composite likelihood ≈ 8087) and version 1.0 (right; negative log composite likelihood ≈ 10051) with 1 hybridization. Hybrid inheritance proportions are labelled in cyan font.

**Figure 17.**
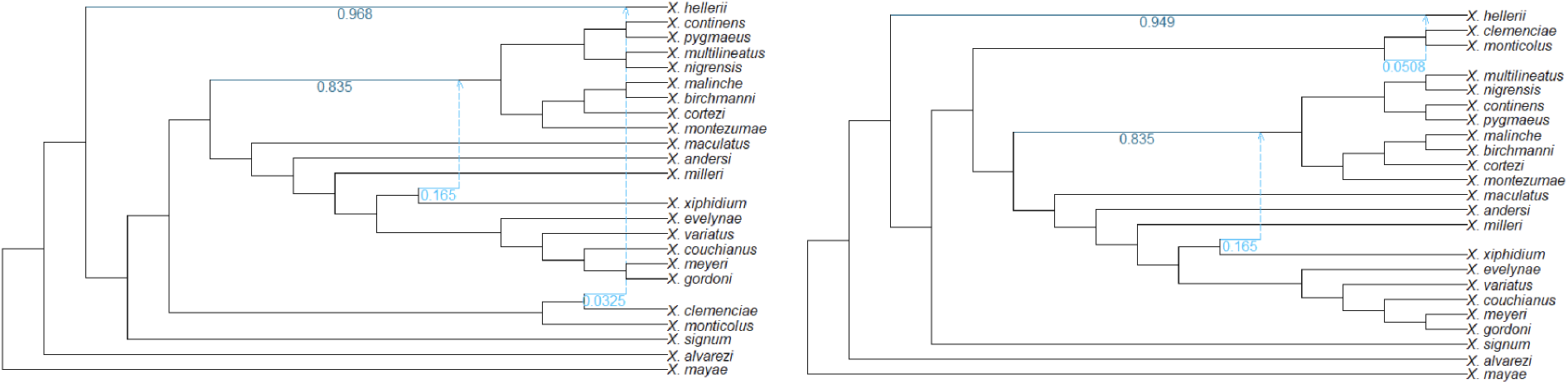
Highest negative log likelihood empirical network inferred by `SNaQ.jl` version 1.1 (left; negative log composite likelihood ≈ 8010) and version 1.0 (right; negative log composite likelihood ≈ 8015) with 2 hybridizations. Hybrid inheritance proportions are labelled in cyan font.

**Figure 18.**
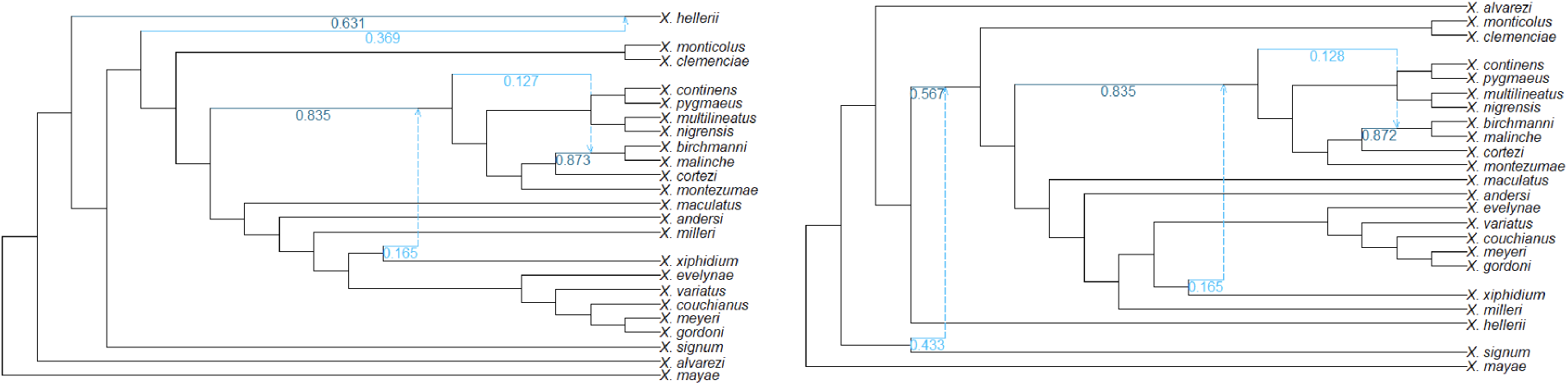
Highest negative log likelihood empirical network inferred by `SNaQ.jl` version 1.1 (left; negative log composite likelihood ≈ 7045) and version 1.0 (right; negative log composite likelihood ≈ 7268) with 3 hybridizations. Hybrid inheritance proportions are labelled in cyan font.

**Figure 19.**
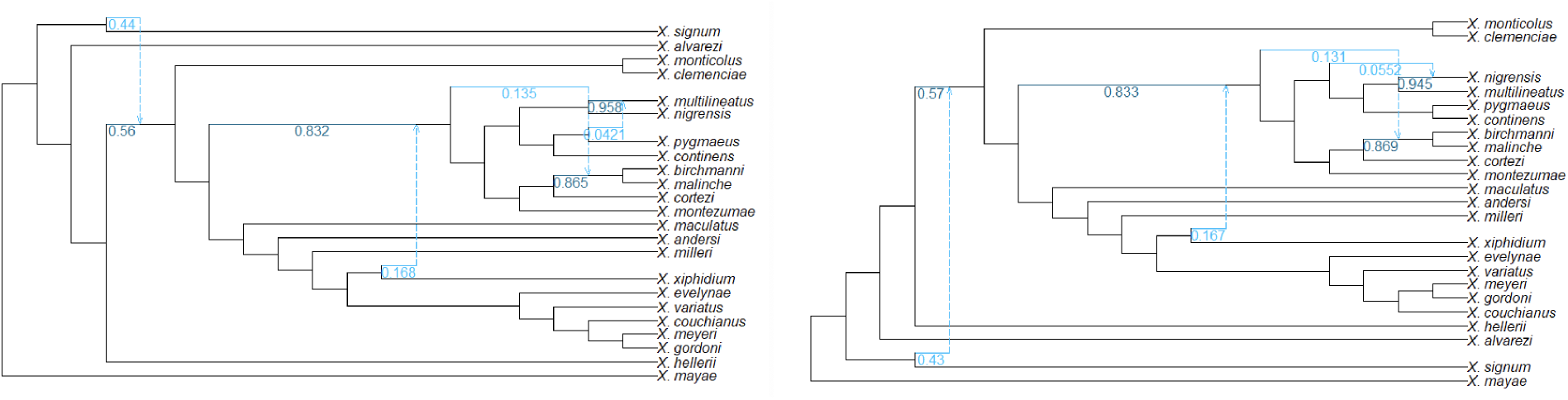
Highest negative log likelihood empirical network inferred by `SNaQ.jl` version 1.1 (left; negative log composite likelihood ≈ 6911) and version 1.0 (right; negative log composite likelihood ≈ 7053) with 4 hybridizations. Hybrid inheritance proportions are labelled in cyan font.

**Figure 20.**
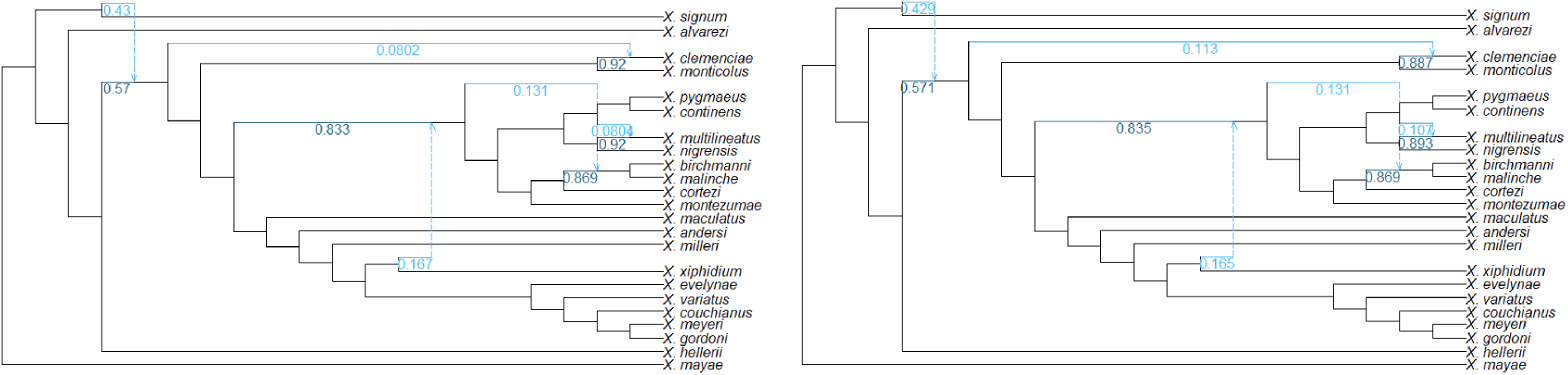
Highest negative log likelihood empirical network inferred by `SNaQ.jl` version 1.1 (left; negative log composite likelihood ≈ 6911) and version 1.0 (right; negative log composite likelihood ≈ 6943) with 5 hybridizations. Hybrid inheritance proportions are labelled in cyan font.

#### A2.2 Model selection curves

**Figure 21.**
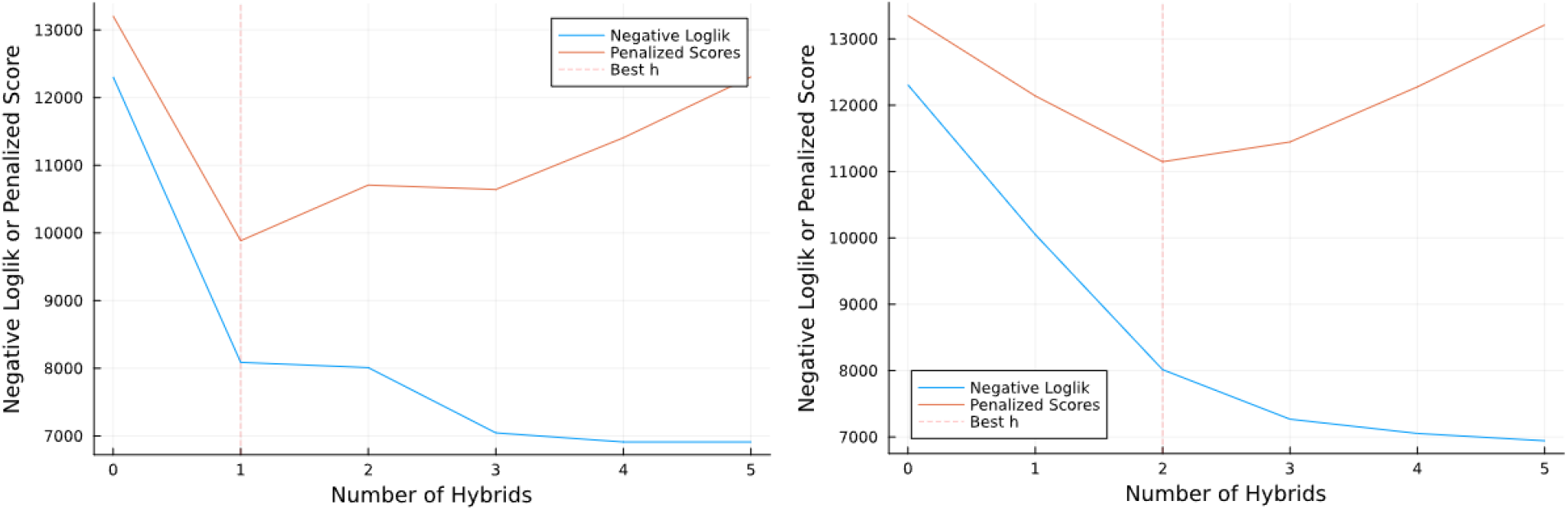
Best inferred log composite likelihoods, slope heuristic penalized scores, and selected model for each empirical network with *h* ∈ {0, 1, 2, 3, 4, 5} inferred by `SNaQ.jl` version 1.1 (left) and version 1.0 (right).

#### A2.3 Networks inferred with *h* = 2 **and** propQuartets *<* 1

**Figure 22.**
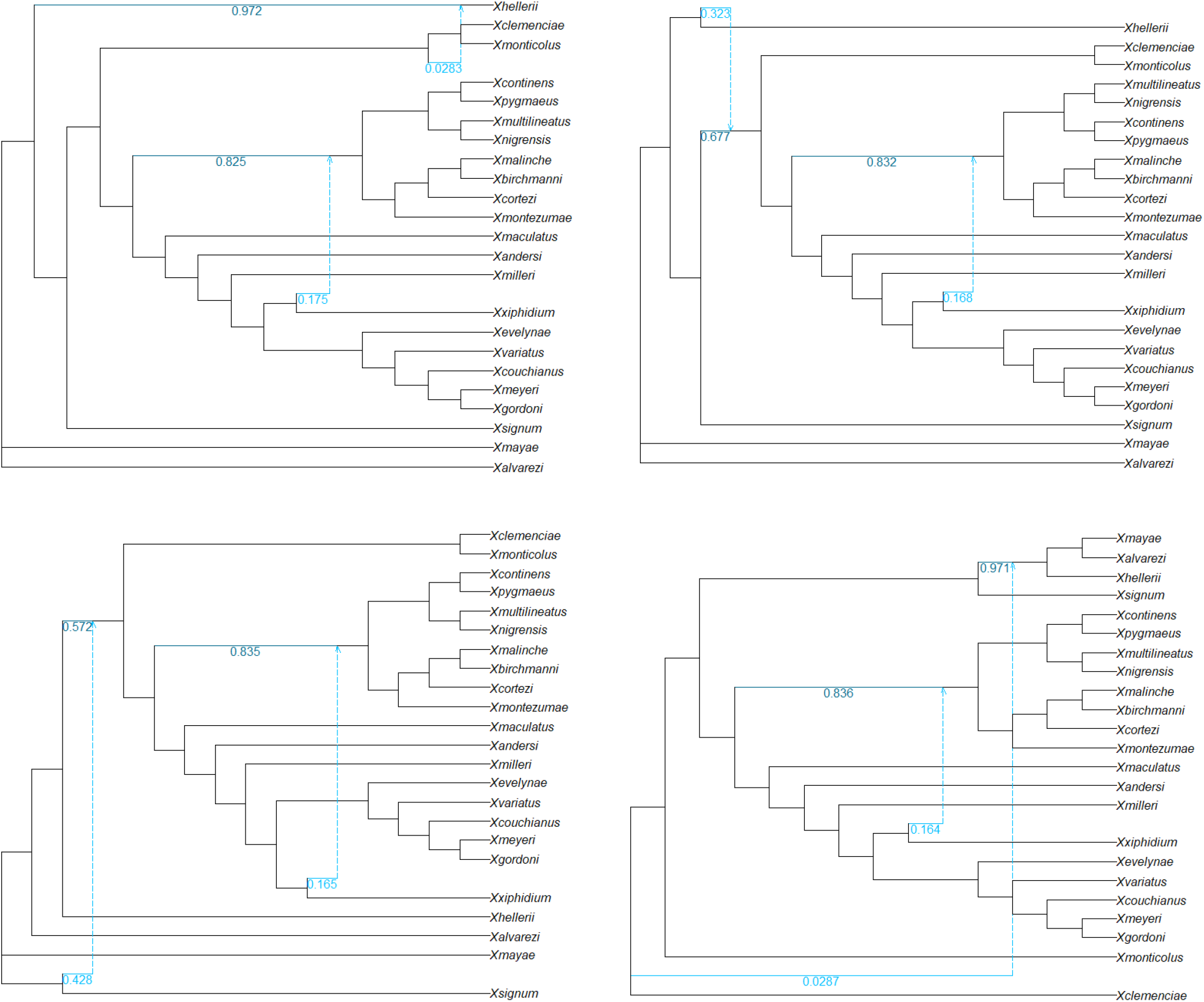
Empirical networks inferred with *h* = 2 and various `propQuartets` values. **Top left:** `propQuartets` = 0.1 (-loglik ≈ 8080), **top right:** `propQuartets` = 0.3 (-loglik ≈ 7729), **bottom left:** `propQuartets` = 0.5 (-loglik ≈ 7663), **bottom right:** `propQuartets` = 0.7 (-loglik ≈ 8020).

## Notes

### Competing Interest Statement

The authors have declared no competing interest.

### Summary of Updates

Several clarifications and lines edits suggested by reviewers. Added an analysis of the empirical network when inferred with propQuartets < 1.0.

## References

J.-P. Baudry, C. Maugis, and B. Michel. Slope heuristics: Overview and implementation. Statistics and Computing, 22(2):455–470, Mar. 2012. ISSN 1573-1375. doi: 10.1007/s11222-011-9236-1.

D. A. Baum. Concordance trees, concordance factors, and the exploration of reticulate genealogy. TAXON, 56(2):417–426, May 2007. ISSN 00400262. doi: 10.1002/tax.562013.

D. Bryant and V. Moulton. Neighbor-net: An agglomerative method for the construction of phylogenetic networks. Molecular Biology and Evolution, 21(2):255–265, 02 2004. ISSN 0737-4038. doi: 10.1093/molbev/msh018. URL https://doi.org/10.1093/molbev/msh018.

J. Chifman and L. Kubatko. Quartet inference from SNP data under the coalescent model. Bioinformatics, 30(23):3317–3324, Dec. 2014. ISSN 1460-2059, 1367-4803. doi: 10.1093/bioinformatics/btu530.

J. Chifman and L. Kubatko. Identifiability of the unrooted species tree topology under the coalescent model with time-reversible substitution processes, site-specific rate variation, and invariable sites. Journal of Theoretical Biology, 374:35–47, June 2015. ISSN 00225193. doi: 10.1016/j.jtbi.2015.03.006.

R. Cui, M. Schumer, K. Kruesi, R. Walter, P. Andolfatto, and G. G. Rosenthal. Phylogenomics reveals extensive reticulate evolution in Xiphophorus fishes: Phylogenomics of Xiphophorus fishes. Evolution, 67(8):2166–2179, Aug. 2013. ISSN 00143820. doi: 10.1111/evo.12099.

J. H. Degnan. Modeling hybridization under the network multispecies coalescent. Systematic Biology, 67(5):786–799, Sept. 2018. ISSN 1063-5157, 1076-836X. doi: 10.1093/sysbio/syy040.

J. Fogg, E. S. Allman, and C. Ané. PhyloCoalSimulations: A Simulator for Network Multispecies Coalescent Models, Including a New Extension for the Inheritance of Gene Flow. Systematic Biology, 72(5):1171–1179, Nov. 2023. ISSN 1063-5157, 1076-836X. doi: 10.1093/sysbio/syad030.

M. Hasegawa, H. Kishino, and T.-a. Yano. Dating of the human-ape splitting by a molecular clock of mitochondrial dna. Journal of Molecular Evolution, 22(2):160–174, Oct. 1985. ISSN 1432-1432. doi: 10.1007/BF02101694.

H. A. Hejase and K. J. Liu. A scalability study of phylogenetic network inference methods using empirical datasets and simulations involving a single reticulation. BMC Bioinformatics, 17(1): 422, Dec. 2016. ISSN 1471-2105. doi: 10.1186/s12859-016-1277-1.

D. H. Huson and D. Bryant. Application of phylogenetic networks in evolutionary studies. Molecular Biology and Evolution, 23(2):254–267, Feb. 2006. ISSN 1537-1719, 0737-4038. doi: 10.1093/molbev/msj030.

J. A. Justison, C. Solis-Lemus, and T. A. Heath. SiPhyNetwork : An R package for simulating phylogenetic networks. Methods in Ecology and Evolution, 14(7):1687–1698, July 2023. ISSN 2041-210X, 2041-210X. doi: 10.1111/2041-210X.14116.

N. Kolbow, S. Kong, and C. Solís-Lemus. A method for massively scalable inference of phylogenetic networks. bioRxiv, May 2025. doi: 10.1101/2025.05.05.652278.

S. Kong, J. C. Pons, L. Kubatko, and K. Wicke. Classes of Explicit Phylogenetic Networks and their Biological and Mathematical Significance. arXiv:2109.10251 [math, q-bio], Jan. 2022a.

S. Kong, D. L. Swofford, and L. S. Kubatko. Inference of Phylogenetic Networks from Sequence Data using Composite Likelihood. Preprint, Evolutionary Biology, Nov. 2022b.

S. Kong, C. Solís-Lemus, and G. P. Tiley. Phylogenetic networks empower biodiversity research. Proceedings of the National Academy of Sciences, 122(31):e2410934122, 2025. doi: 10.1073/pnas.2410934122. URL https://www.pnas.org/doi/abs/10.1073/pnas.2410934122.

L. Liu, L. Yu, and S. V. Edwards. A maximum pseudo-likelihood approach for estimating species trees under the coalescent model. BMC Evolutionary Biology, 10(1):302, 2010. ISSN 1471-2148. doi: 10.1186/1471-2148-10-302.

L.-T. Nguyen, H. A. Schmidt, A. von Haeseler, and B. Q. Minh. IQ-TREE: A fast and effective stochastic algorithm for estimating maximum-likelihood phylogenies. Molecular Biology and Evolution, 32(1):268–274, Jan. 2015. ISSN 1537-1719, 0737-4038. doi: 10.1093/molbev/msu300.

One Thousand Plant Transcriptomes Initiative. One thousand plant transcriptomes and the phylogenomics of green plants. Nature, 574(7780):679–685, 2019.

A. Rambaut and N. C. Grass. Seq-Gen: An application for the Monte Carlo simulation of DNA sequence evolution along phylogenetic trees. Bioinformatics, 13(3):235–238, 1997. ISSN 1367-4803, 1460-2059. doi: 10.1093/bioinformatics/13.3.235.

L. J. Revell. Phytools: An R package for phylogenetic comparative biology (and other things). Methods in Ecology and Evolution, 3(2):217–223, Apr. 2012. ISSN 2041-210X, 2041-210X. doi: 10.1111/j.2041-210X.2011.00169.x.

U. Rosas-Puchuri, N. Kolbow, C. Solís-Lemus, S. Khanmohammadi, and B.-R. Ricardo. Sparse learning for scalable phylogenetic network inference, Nov. 2025. ISSN 2692-8205.

C. Solís-Lemus and C. Ané. Inferring phylogenetic networks with maximum pseudolikelihood under incomplete lineage sorting. PLOS Genetics, 12(3):e1005896, Mar. 2016. ISSN 1553-7404. doi: 10.1371/journal.pgen.1005896.

C. Solís-Lemus, P. Bastide, and C. Ané. PhyloNetworks: A package for phylogenetic networks. Molecular Biology and Evolution, 34(12):3292–3298, Dec. 2017. ISSN 0737-4038, 1537-1719. doi: 10.1093/molbev/msx235.

D. Wen, Y. Yu, and L. Nakhleh. Bayesian inference of reticulate phylogenies under the multispecies network coalescent. PLOS Genetics, 12(5):e1006006, May 2016. ISSN 1553-7404. doi: 10.1371/journal.pgen.1006006.

D. Wen, Y. Yu, J. Zhu, and L. Nakhleh. Inferring phylogenetic networks using phylonet. Systematic Biology, 67(4):735–740, July 2018. ISSN 1063-5157, 1076-836X. doi: 10.1093/sysbio/syy015.

Y. Yu and L. Nakhleh. A maximum pseudo-likelihood approach for phylogenetic networks. BMC Genomics, 16(S10):S10, Dec. 2015. ISSN 1471-2164. doi: 10.1186/1471-2164-16-S10-S10.

Y. Yu, J. H. Degnan, and L. Nakhleh. The probability of a gene tree topology within a phylogenetic network with applications to hybridization detection. PLoS Genetics, 8(4):e1002660, Apr. 2012. ISSN 1553-7404. doi: 10.1371/journal.pgen.1002660.

Y. Yu, J. Dong, K. J. Liu, and L. Nakhleh. Maximum likelihood inference of reticulate evolutionary histories. Proceedings of the National Academy of Sciences, 111(46):16448–16453, Nov. 2014. ISSN 0027-8424, 1091-6490. doi: 10.1073/pnas.1407950111.

C. Zhang and S. Mirarab. Weighting by Gene Tree Uncertainty Improves Accuracy of Quartet-based Species Trees. Molecular Biology and Evolution, 39(12):msac215, Dec. 2022. ISSN 0737-4038, 1537-1719. doi: 10.1093/molbev/msac215.

C. Zhang, H. A. Ogilvie, A. J. Drummond, and T. Stadler. Bayesian inference of species networks from multilocus sequence data. Molecular Biology and Evolution, 35(2):504–517, Feb. 2018. ISSN 0737-4038, 1537-1719. doi: 10.1093/molbev/msx307.

J. Zhu and L. Nakhleh. Inference of species phylogenies from bi-allelic markers using pseudo-likelihood. Bioinformatics, 34(13):i376–i385, July 2018. ISSN 1367-4803, 1460-2059. doi: 10.1093/bioinformatics/bty295.

